# Evolutionary dynamics of roX lncRNA function and genomic occupancy

**DOI:** 10.1101/026013

**Authors:** Jeffrey J. Quinn, Qiangfeng C. Zhang, Plamen Georgiev, Ibrahim A. Ilik, Asifa Akhtar, Howard Y. Chang

## Abstract

Many long noncoding RNAs (lncRNAs) can regulate chromatin states, but the evolutionary origin and dynamics driving lncRNA–genome interactions are unclear. We developed an integrative strategy that identifies lncRNA orthologs in different species despite limited sequence similarity that is applicable to fly and mammalian lncRNAs. Analysis of the roX lncRNAs, which are essential for dosage compensation of the single X-chromosome in Drosophila males, revealed 47 new roX orthologs in diverse Drosophilid species across ∼40 million years of evolution. Genetic rescue by roX orthologs and engineered synthetic lncRNAs showed that evolutionary maintenance of focal structural repeats mediates roX function. Genomic occupancy maps of roX RNAs in four species revealed rapid turnover of individual binding sites but conservation within nearby chromosomal neighborhoods. Many new roX binding sites evolved from DNA encoding a pre-existing RNA splicing signal, effectively linking dosage compensation to transcribed genes. Thus, evolutionary analysis illuminates the principles for the birth and death of lncRNAs and their genomic targets.

---

Eukaryotic genomes are replete with long noncoding RNA (lncRNA) genes that are diverse, tightly regulated, and engaged in numerous biological processes^1-4^. LncRNAs differ from protein-coding genes in many ways^5^ and in particular, lncRNAs are typically less conserved at the level of primary sequence^6, 7^. This is because many of the selection pressures that constrain protein-coding primary sequences do not apply to lncRNAs, such as maintenance of open reading frames and codon synonymy. The low primary sequence conservation has led some to dismiss lncRNAs as transcriptional noise^8^, and the discovery of lncRNA orthologs in other genomes is challenging. These issues in turn limit the investigation of lncRNAs’ evolutionary origins and dynamics, conserved elements, and functions. Examples of such evolutionary analyses are scarce yet valuable, like the study that revealed the independent evolutionary origins of the mammalian dosage compensation lncRNAs Xist and Rsx, with Xist having arisen from a pseudogenized protein-coding gene^9, 10^. Despite many predictions from RNA-seq data^11, 12^, few lncRNA orthologs that function across species have been experimentally verified.

Besides their primary sequence, other lncRNA features are better conserved: including synteny, short sequence homology, and secondary structure^6, 7, 11, 13^. LncRNA synteny – the order and orientation of neighboring genes – may be conserved because many lncRNAs act in *cis*^14^, are *cis*-defined (e.g. divergent transcripts^15^), serve as chromosome landmarks (e.g. XIST at the X-inactivation center^9^), or simply because intra-chromosomal gene shuffling often preserves genomic blocks^16, 17^. Short sequences conservation (i.e. “microhomology”) may result from tight sequence constraint of functional elements, like the microRNA binding site on cyrano lncRNA^6^, and can be identified by linear alignment or motif discovery tools. In some cases, microhomologous elements are repeated within a lncRNA and can act as protein binding sites, such as Xist Repeats A and C^18-20^ or the roXboxes of roX lncRNAs^21^. Secondary structures of lncRNAs may be conserved if they confer specific functions, mirroring the structure-function relationship of proteins, like the RNA triplex and tRNA-like cloverleaf structures of MALAT1 lncRNA^22^.

Two lncRNAs in *Drosophila melanogaster*, roX1 and roX2, are essential for dosage compensation, wherein gene expression from the single X-chromosome in males is doubled to match gene expression of females’ two X-chromosomes. roX lncRNAs are critical for assembling the dosage compensation ribonucleoprotein complex (DCC), targeting the DCC to hundreds of high-affinity sites (HAS) on the X-chromosome, and spreading the DCC to epigenetically activate genes^23-25^. Genetic ablation of both *roX* genes or any of five DCC proteins results in failed dosage compensation and male-specific lethality^26^. Despite the fact that roX1 and roX2 are functionally redundant, they differ greatly in size (3.7kb and 0.6kb, respectively), sequence, and expression patterns. The functional redundancy between roX RNAs is primarily attributed to a short, repeated sequence motif (the 8-nucleotide roXbox motif) embedded in stem-loop structures in both roX1 and roX2^21, 27^. Previous roX ortholog search efforts identified eight roX1 and nine roX2 orthologs in other species using whole-gene BLAST^28, 29^ and structure detection by sequence covariation^30^ (indicated in **Figure 1B**); however, these queries failed to identify roX orthologs in many Drosophila species, as the primary sequence identity between discovered orthologs was close to random^28^. Nonetheless, these studies highlighted evolutionarily conserved structures that are essential to roX function^21, 27, 31^.

**Figure 1.**
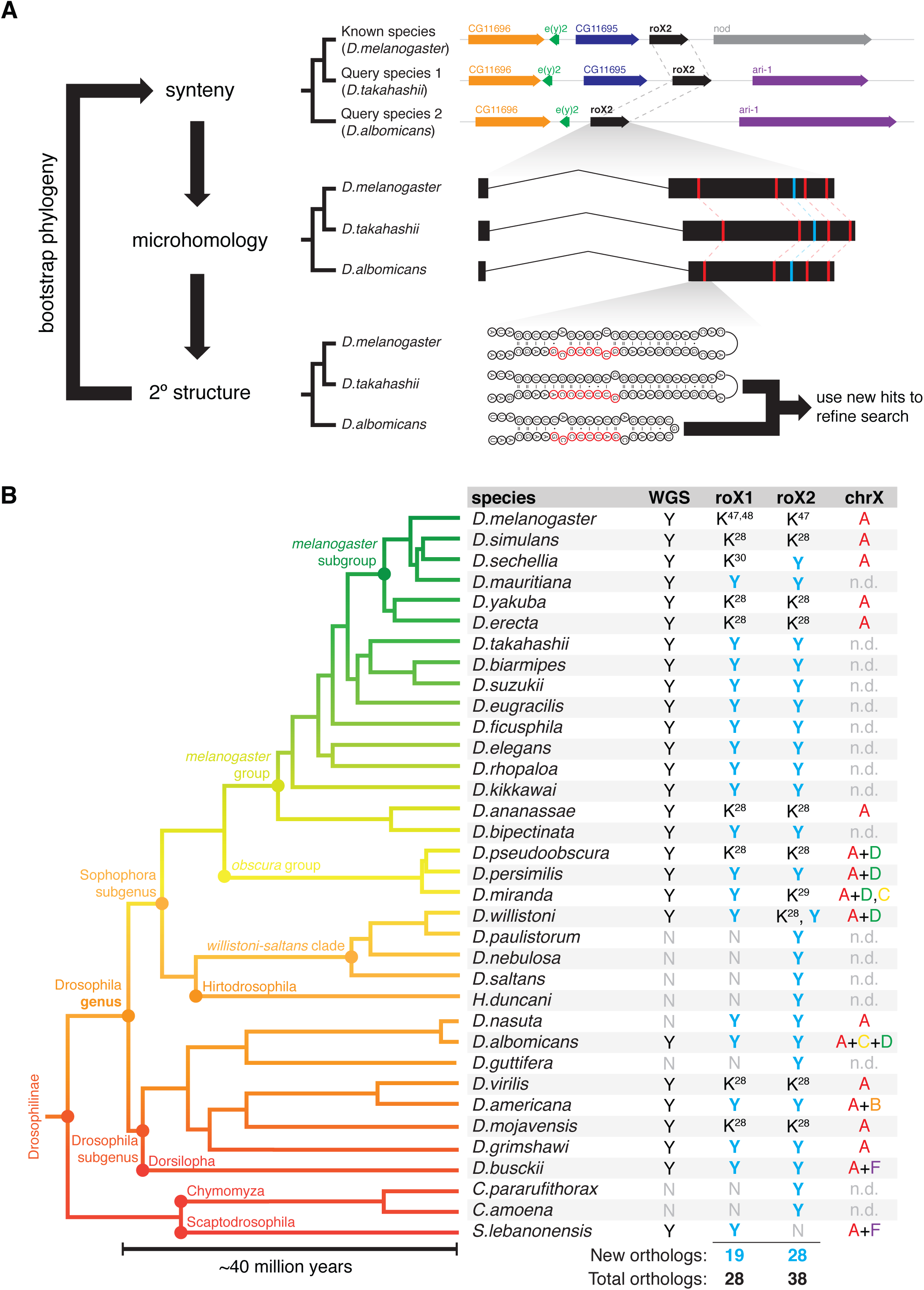
Summary of lncRNA ortholog search strategy and queried species. **(A)** The search strategy finds lncRNA orthologs in query species by integrating synteny, microhomology, and secondary structure features of a known lncRNA. The search features are iteratively refined by bootstrapping new lncRNA candidate hits and the phylogenetic relationships between queried species. A priori knowledge of only a single lncRNA is needed to initiate the search. **(B)** Phylogenetic tree of the 35 Drosophilid species queried in this study. Whole genome sequencing (WGS) assemblies were available for 27 species. Nine roX1 and ten roX2 orthologs have previously been described (K, known roX ortholog from refs. ^28-30, 48, 49^); our search identified 47 new roX orthologs (Y, new ortholog; N, no ortholog found). X-chromosome karyotypes are indicated by Müller elements (n.d., no data).

Here, we describe a lncRNA ortholog search strategy that integrates synteny, microhomology, and secondary structure (**Figure 1A**). Application of this strategy to roX lncRNAs in 35 diverse fruitfly species discovered 47 novel roX orthologs. Genome-wide roX RNA occupancy maps in four species revealed distinct evolutionary principles that shape lncRNA structure, function, and genomic binding sites.

## Nested homology: lncRNA ortholog search strategy

The Drosophila genus is highly diverse, comprising nearly 2000 named species that diverged ∼40 Mya (Sophophora-Drosophila subgenera divergence^32, 33^) with well-defined phylogenetic relations^34^ (**Figure 1B**). For this study, we selected 27 flies with sequenced genomes plus 8 additional species to maximize phylogenetic diversity, including the outgroup genera Chymomyza and Scaptodrosophila (*summarized in* **Figure 1B**).

Our lncRNA ortholog search strategy consists of three main steps (synteny, microhomology, and secondary structure) and iteratively bootstraps new ortholog hits and known phylogenetic relationships (**Figure 1A**). First, we searched for synteny blocks likely containing the *roX1* or *roX2* loci, employing a computational or analog method (tBLASTn or degenerate PCR, respectively), depending on the availability of completed genome assemblies for the subject species. Next, we homed in on roX orthologs by searching for incidences of microhomology (roXbox motifs) and structure (roXbox stem-loops) within the identified synteny block, which thus served as landmarks for the roX ortholog candidates. Lastly, we leveraged new lncRNA ortholog hits and the defined phylogenetic relationships between species to iteratively refine the search parameters. For example, we collapsed roXboxes from each newly identified roX ortholog to further refine the motif; we also searched synteny blocks matched to the closest relative and its *roX* locus. In this way, the search strategy became more powerful with each new ortholog identified.

## Search strategy yielded 47 new roX orthologs

This search uncovered 47 new roX ortholog candidates (19 roX1s and 28 roX2s) in addition to those previously described, more than tripling the number of known roX orthologs (66 total; **Figure 1B**). In the few cases where roX orthologs could not be identified, a complete genomic assembly was lacking or incomplete or there was syntenic disruption at the *roX* locus. Curiously, the search identified three high-scoring roX homolog candidates in *D.willistoni*; close analysis of these candidates in *D.willistoni* and its relatives indicated that *roX2* was duplicated in the *willistoni-saltans* clade after the divergence of *H.duncani*, resulting in up to three functional *roX* genes (we call this *roX2* paralog “*roX3*”; **figure supplement 1**). The roX orthologs identified exhibit exceptionally low primary sequence conservation, dropping to the lower limit of homology (scrambled sequences) when comparing sequences between Sophophora and Drosophila subgenera or outgroups (**Figure 2A-B**).

**Figure 2.**
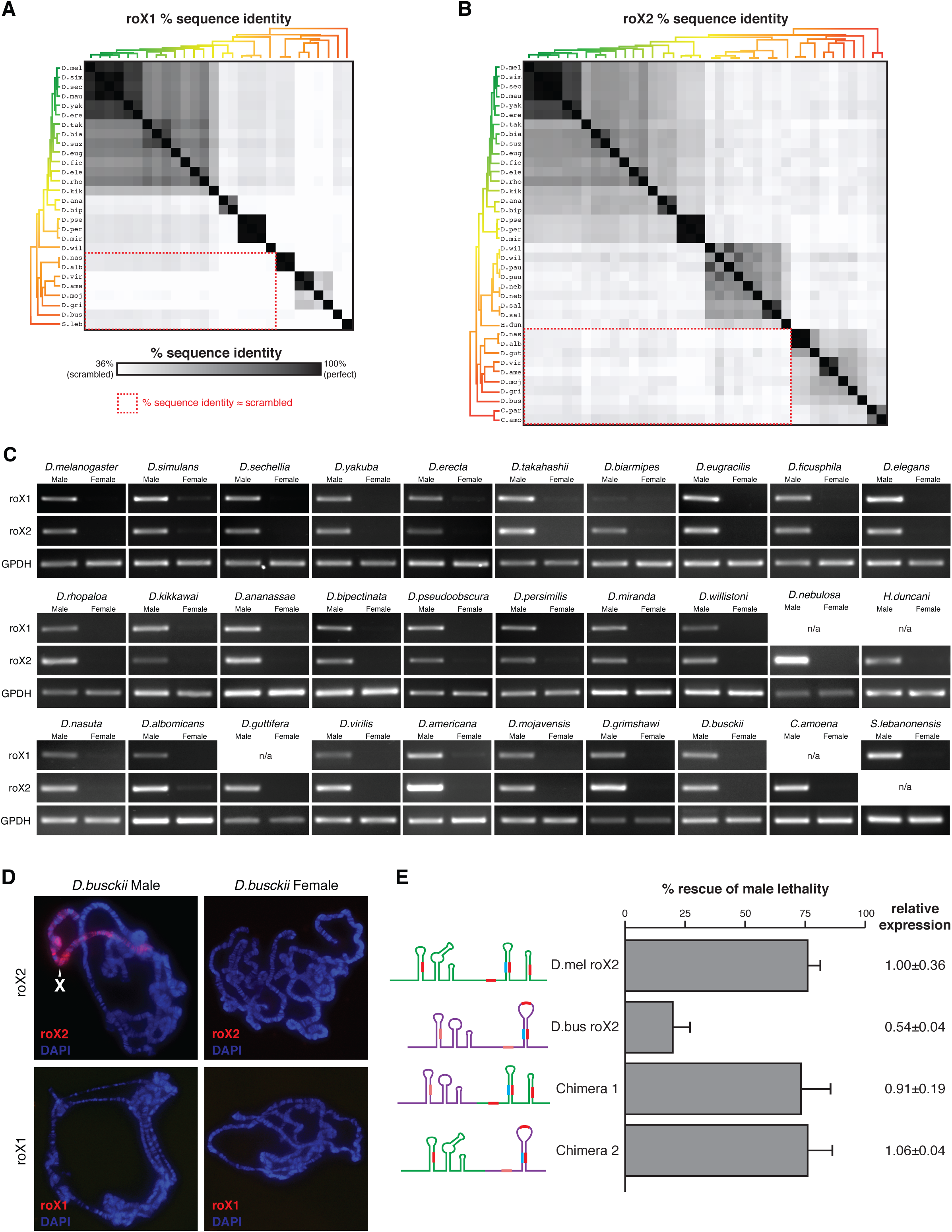
Figure 2. Identified roX candidates are bona fide orthologs despite sequence divergence. (**A, B**) Heatmap showing the sequence conservation between the identified roX1 (**A**) and roX2 (**B**) ortholog candidates relative to the lower limit of homology (scrambled sequences). Phylogenetic trees as in **Figure 1B**. Red dashed boxes highlight exceptionally poor conservation between distantly related species. **(C)** RT-PCR of roX1, roX2, and GPDH RNA in male and female flies. roX1 and roX2 orthologs exhibit strong male-biased expression; GPDH is a sex-independent control (n/a, no ortholog found). **(D)** RNA FISH of roX1 and roX2 in polytene chromosomes from male and female *D.busckii* larvae. roX2 paints the male X-chromosome (white arrowhead), but not the female X; roX1 is not detected. **(E)** Rescue of male lethality in *roX*-null *D.melanogaster* (D.mel) males *D.busckii* (D.bus) roX2 or chimeric *busckii – melanogaster* roX2 transgenes. RNA cartoons depict secondary structures, with roXboxes and inverted roXboxes as filled red and cyan rectangles, respectively. Error bars show standard deviation. Expression is calculated relative to wild-type roX2 transgene, ±standard deviation.

Additionally, to test the generalizability of this search strategy, we searched for orthologs of the HOTAIR lncRNA in vertebrate genomes, initiating the search with only the sequence of human HOTAIR in the *HOXC* cluster. We identified the orthologous *HOTAIR* locus in 33 eutherian genomes, and evidence for conservation of numerous sequence elements in genomes as evolutionarily deep as coelacanth and zebrafish (**figure supplement 15**). The putative HOTAIR orthologs are encoded within the same genomic locus (between *HOXC11* and *HOXC12*) and have short conserved sequence elements; several experimentally verified RNA structures in HOTAIR^35^ show signatures of evolutionary conservation.

## Ortholog candidates are bona fide roX genes

Male-biased expression, RNA localization, and genetic rescue confirmed that the identified roX candidates were bona fide roX orthologs. We first assayed their expression in whole adult males and females by RT-PCR using species-specific primers for the roX ortholog candidates and a housekeeping mRNA (GPDH). In all 30 species tested, the roX1 and roX2 candidates were transcribed RNAs and displayed strong male-biased expression (**Figure 2C**).

We next used RNA FISH (fluorescence *in situ* hybridization) to investigate the localization of roX1 and roX2 on *D.busckii* polytene chromosome squashes. *D.busckii* was selected because of its basal position within the Drosophila subgenus, substantial evolutionary distance from *D.melanogaster* (diverged ∼40 Mya^32, 33^), and low homology with other roX orthologs (**Figure 2B**). Notably, *D.busckii* roX2 paints the X-chromosome in males but not females, and roX1 was not detected in either (**Figure 2D**). This localization pattern matches that of *D.mojavensis* and *D.virilis* (both also in the Drosophila subgenus), in which roX2 – but not roX1 – coats the male X-chromosome^28^.

Next we asked if transgenic expression of *D.busckii* roX2 could rescue male lethality in *roX*-null *D.melanogaster*. As positive control, transgenic expression of *D.melanogaster* roX2 rescued ∼75% of males (**Figure 2E**). Notably, *D.busckii* roX2 rescued ∼20% of males, which – though modest – is substantially greater than *roX*- null background (<0.01% male viability^36^). Complete structural disruption of the 5’- and 3’-halves of *D.melanogaster* roX2 abrogates male rescue^21^, but two chimeric fusions of *melanogaster–busckii* roX2 halves rescued males as robustly as the positive control (**Figure 2E**). Prior work showed that *roX*-null *D.melanogaster* males are best rescued by roX transgenes from *D.melanogaster*, followed by *D.ananassae*, then *D.willistoni*, suggesting that rescue efficiency decreases with increasing evolutionary distance^27^. The rescue by *D.busckii* roX2 fits this trend, and confirms it as a bona fide roX2 ortholog. Because our bootstrapping strategy uses a chain of roX orthologs to iteratively bridge distantly related species, successful rescue by *D.busckii* roX2 implies that the intervening roX2 candidates are true orthologs as well. In addition, enhanced rescue by the chimeric RNAs demonstrates the modular nature of structured repeats in lncRNAs^37^.

## Conserved features of lncRNA genes

Given that our search strategy begins by analyzing synteny, it is not surprising that most roX orthologs identified had conserved gene neighbors (**figure supplement 2**). In *D.melanogaster*, *roX1* and *roX2* loci are on the X-chromosome; and in all other species, the neighboring genes are also X-linked, suggesting that *roX* orthologs are similarly X-linked. This mirrors the finding that *Xist* orthologs in eutherians are always encoded on the X-chromosome^16^. Using 5’- and 3’-RACE, we showed that roX2 orthologs share a similar exon–intron gene structure, alternative splicing and polyadenylation pattern, and gene length (**figure supplement 3**). roX2 roXboxes are the most prominently conserved sequences in primary sequence, relative position, and orientation.

We found conserved structures in roX1 and roX2, some previously described to be functional^21, 27, 28^, as well as many novel ones (**figure supplements 4, 5**). For example, roX1-D3 domain contains a highly conserved stem-loop (IRB–RB) that was ultraconserved in every roX1 ortholog found (**figure supplement 4C**). Interestingly, another stem-loop in roX1-D3 is only present in the Sophophora subgenus and *S.lebanonensis*, but is absent in the Drosophila subgenus (**figure supplement 4B**), despite being a primary binding site for the DCC and important for roX1 function in *D.melanogaster*^21, 37^. Similarly, structures within roX1-D2 are lost in *D.willistoni* (**figure supplement 4D**). The absence of such important structures in these species may have consequences for roX1 function. Their presence in the outgroup species *S.lebanonensis* indicates that they were lost in the Drosophila subgenus (rather than gained in the Sophophora subgenus). We also found evidence for an ultraconserved structure (**figure supplement 5B**) in roX2, as well as complex structures wherein two or more roXboxes compete for one intervening inverted roXbox, indicative of alternative secondary structures (RB4–IRB and IRB–RB5; **figure supplement 5C**). These structures are arranged on roX2 exon-3 in a similar configuration in all species (**figure supplement 6**).

## roX orthologs bind the X-chromosome

The fruitfly genome consists of six chromosome arms, called Müller elements (ME)-A–F; the X-chromosome in *D.melanogaster* is ME-A. However, the X-chromosome in flies has undergone numerous karyotype reversals and ME fusions throughout evolution^38^, such as the ME-A+D fusion in *D.willistoni* (**Figure 1B**). Previous studies have found that newly evolved sex chromosomes can rapidly acquire DCC binding sites, through amplification of simple GA-dinucleotide repeats that approximate the MSL recognition element (MRE) or domestication of MRE-bearing transposable elements^29, 39, 40^.

To understand the evolution of lncRNA–genome interactions, we mapped the genomic binding sites of roX1 and roX2 orthologs in four species: *D.melanogaster, D.willistoni, D.virilis,* and *D.busckii*. We chose these four species as representatives for the Drosophila genus’ diversity and distinct X-chromosome karyotypes (**Figure 1B**). We developed methods to perform *in vivo* ChIRP-seq (Chromatin Isolation by RNA Purification and sequencing) directly from homogenized whole larvae. In ChIRP-seq, chromatin is cross-linked and fragmented; the target RNA and associated chromatin are affinity-purified with biotinylated antisense oligonucleotide probes; next, the co-purified DNA is sequenced (**Figure 3A**). Thus, ChIRP-seq maps the *in vivo* genomic binding sites of a chromatin-associated RNA from endogenous interactions^24^. Unlike ChIP-seq (chromatin immunoprecipitation) in diverse species, which may require species-specific antibodies or transgenic epitope-tagging systems, ChIRP-seq in diverse species requires only new antisense oligonucleotide sequences that can be readily designed from lncRNA sequences, regardless of how divergent they may be.

**Figure 3.**
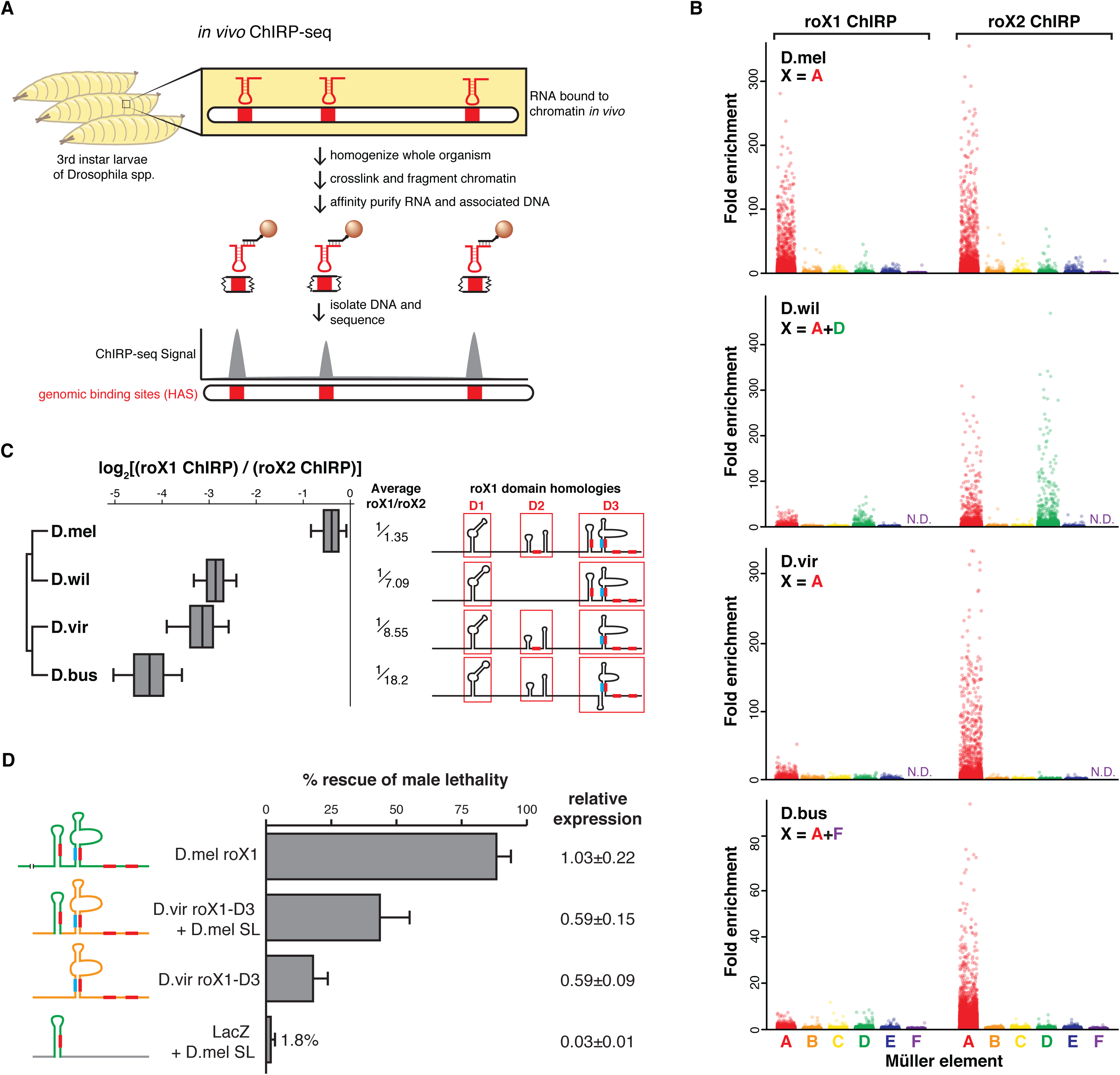
Genomic occupancy maps of roX orthologs highlight the loss of roX1 – roX2 functional redundancy. **(A)** ChIRP-seq identifies the genome-wide binding sites of an RNA target, performed directly from chromatin prepared from Drosophila larvae. **(B)** roX1 and roX2 signal enrichment (ChIRP/input) in 1kb windows of Müller elements A-F in four Drosophila species. Signal is enriched on the X-chromosome. roX1 enrichment is diminished relative to roX2 in *D.willistoni* (D.wil), *D.virilis* (D.vir), and *D.busckii*. (N.D., no data, as no scaffolds mapped to ME-F.) (**C**, **left**) The ratio of roX1 to roX2 ChIRP signal at binding sites shows that roX2 is the dominant roX RNA in *D.willistoni*, *D.virilis*, and *D.busckii*. Average roX1/roX2 bias is shown as fraction. (**C, right**) Known functional domains (red outlines), secondary structures, and roXboxes (filled red or cyan rectangles) of roX1 are absent in *D.willistoni*, *D.virilis*, and *D.busckii*. Only *D.melanogaster* roX1 has a full complement of these repetitive elements. See also **figure supplement 4**. (**D**) Rescue of male lethality in *roX*-null *D.melanogaster* males improves with the number of repetitive roX elements. LacZ with *D.melanogaster* stem-loop (SL) rescues poorly, *D.virilis* roX1-D3 rescues modestly, and addition of the *D.melanogaster* stem-loop to *D.virilis* roX1-D3 further improves rescue, approaching the wild-type *D.melanogaster* roX1 rescue efficiency. Error bars and relative expression as in **Figure 2E**.

We performed roX1 and roX2 ChIRP-seq in the four species, and mapped the reads to their respective genomes. We assigned scaffolds from each genome assembly to specific MEs based on coding sequence homology to *D.melanogaster* proteins, as done previously^38^, and then calculated ChIRP signal enrichment (ChIRP/input) for each ME in 1kb windows (**Figure 3B**). We found that roX2 preferentially occupied the X-chromosome in each species, including ME-D in *D.willistoni*. Interestingly, the tiny X-fused ME-F was not enriched in *D.busckii*, though this may be the result of the epigenetic silencing of ME-F and incomplete decay of the Y-fused ME-F^38, 41^. The extensive roX2 binding on *D.willistoni* ME-D further supports the hypothesis that new X-chromosomes evolve novel binding sites rather than modify or exchange DCC components^29^.

Analysis of roX genomic occupancy indicated that roX1–roX2 functional redundancy has degenerated in some species. roX1 and roX2 ChIRP-seq are highly correlated for all species, indicating that within each species roX1 and roX2 bind the same loci, though with unequal potency (**figure supplement 7**). As expected, *D.melanogaster* roX1 and roX2 ChIRP-seq enriched for the X-chromosome to approximately the same extent, but roX1 enrichment showed quantitative differences in the other species (**Figure 3B-C**), despite equivalently effective capture of roX1 and roX2 RNAs in each species (not shown). roX1 enrichment was 7.09-, 8.55-, and 18.2-fold weaker than roX2 in *D.willistoni*, *D.virilis*, and *D.busckii*, respectively (**Figure 3C**). This is consistent with roX1’s apparent absence on the X by RNA FISH in *D.virilis* and *D.busckii*^28^ (**Figure 2D**). The decreasing potency of roX1 in these species is correlated with repeated loss of stem-loops and roXboxes in domains D2 and D3 (**Figure 3C** and **figure supplement 4**).

We tested the functional consequence of the loss of such repetitive structural elements in roX1-D3 domain, using transgenic rescue of *roX*-null *D.melanogaster* males (**Figure 3D**). A transgene containing a single roXbox stem-loop from *D.melanogaster* roX1 embedded in bacterial LacZ mRNA rescued males poorly (1.8%). Although seemingly low, this level of rescue is ∼100-fold improved over *roX*-null flies (<0.01% male viability^36^), and thus such a stem-loop would confer a major selective advantage. Next, wild-type *D.virilis* roX1-D3 modestly rescues males (18%), consistent with its limited repertoire of roXbox stem-loops and modest X-chromosome occupancy. Adding the *D.melanogaster* stem-loop to *D.virilis* roX1-D3 substantially improved male rescue (43%), approaching the rescue by the positive control, *D.melanogaster* roX1 (88%), which rescues to the same extent as roX1-D3 alone^37^. These findings suggest that in the Drosophila subgenus roX1 has vestigial functional importance due to repetitive structural element losses; in flies like *D.melanogaster*, the observed roX1–roX2 functional redundancy results from the retention of such elements.

## Evolution of high-affinity sites

The high-resolution maps of roX RNA binding allowed us to trace the conservation of roX interaction with genomes at the level of chromosomes, genes, and individual DNA elements. High-affinity sites (HAS) are defined by joint binding of roX RNAs, MLE, and MSL2 (DCC proteins which directly bind roX^21, 24, 42^). Close inspection of homologous genomic windows in the four species revealed that the position of most roX-occupied HAS are evolutionarily dynamic (**Figure 4A**), whereas a minority of HAS are at the same location in all species (**Figure 4B**). HAS have conserved characteristics in each species. For example, there are hundreds of HAS on the X-chromosome in each species, and *D.willistoni* has nearly twice as many HAS in accord with its larger X-chromosome (**Figure 4C**). The two HAS within the *roX1* and *roX2* loci were among the strongest binding sites and occupied by both roX RNAs in all species (not shown), consistent with our previous report^37^. The few binding sites found on autosomes in *D.melanogaster* are reproducible and some are conserved in other species^37^ (**figure supplement 8**). In all species, the top enriched DNA motif was a GA-rich DNA sequence located at HAS centers (**Figure 4D** and **figure supplement 9**), matching the MRE motif in *D.melanogaster*. On *D.willistoni*’s ME-D, we do not find enrichment of any other sequences that would support alternative mechanisms of MRE accumulation (**figure supplement 10**); thus the transposable element-taming mechanism observed in *D.miranda* may be unique to *D.miranda* or species with more recently evolved neo-sex karyotypes^40^.

**Figure 4.**
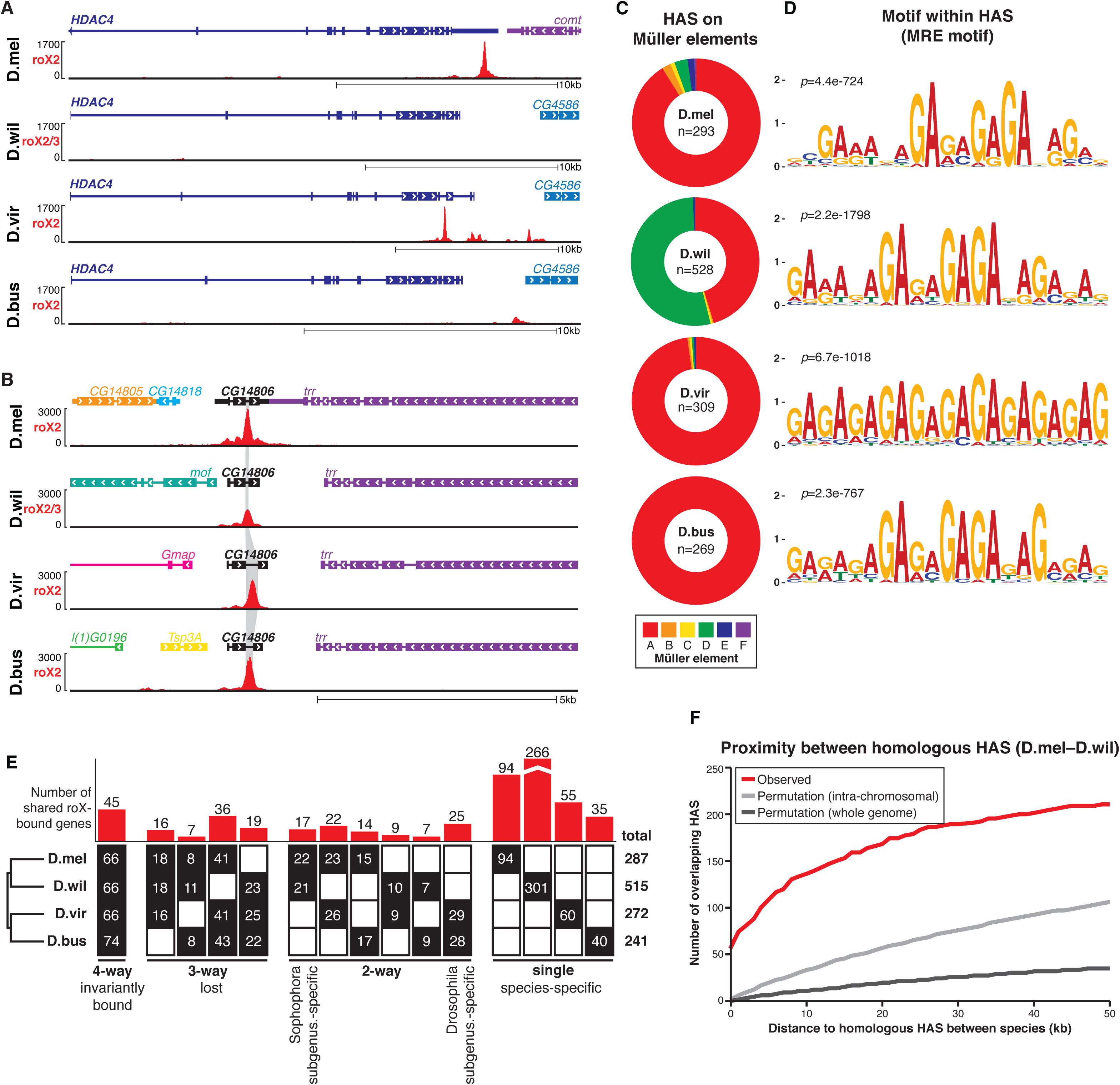
Evolutionary dynamics of roX-bound high-affinity sites (HAS). (**A, B**) Representative genomic windows showing roX2 ChIRP-seq tracks. (**A**) Some HAS are evolutionarily dynamic. The strong HAS in the 3’UTR of *D.melanogaster HDAC4* is absent or present elsewhere in other species. (**B**) Other HAS are evolutionarily conserved, such as the HAS (gray highlight) in the last intron of *CG17841*. **(C)** roX2 ChIRP-seq identified hundreds of HAS on the X-chromosome of each species (*n*, total number of HAS). **(D)** HAS contain the GA-dinculeotide repeat MRE motif. **(E)** Gene-level conservation of HAS between four species. 45 genes are roX-bound (overlapping or neighboring a HAS) in all four species. *D.willistoni* has the most species-specific roX-bound genes because of its larger X-chromosome. Above, number of shared roX-bound genes; below, number of HAS within shared genes in each species. **(F)** Pairwise proximity of orthologous HAS between *D.melanogaster* and *D.willistoni*. ∼60 HAS directly overlap in genomic lift-over (distance=0); however, if an exact HAS homolog is lost (distance>0), another HAS is likely nearby. There are more overlapping HAS than expected by random permutation of HAS within each chromosome or over the whole genome. See also **figure supplement 9**.

Detailed evolutionary analyses revealed that HAS are under selection for proximity, but not precise location relative to genes. We counted the number of inter-species overlapping HAS at the level of genes or DNA elements. At the level of HAS-associated genes (defined as the nearest gene within 1kb of a HAS), we found a small proportion of overlap among the four species (invariantly bound genes; **Figure 4E**). Species-specific HAS-associated genes are the most abundant class (**Figure 4E**, right), indicating poor conservation of the precise genes to which the DCC is targeted. Analysis of the distance between each HAS from one species and the nearest HAS in another species showed that HAS are significantly more likely to directly overlap or be present in the same chromosomal neighborhood than expected by chance alone (from randomly permuting HAS over their respective chromosomes or the whole genome, **Figure 4F** and **figure supplement 11**). The observed species-to-species distance between nearest homologous HAS is most enriched in local genomic neighborhoods up to ∼30kb, and then saturates. Thus, HAS exhibit a conservation pattern that is similar to transcriptional enhancers^43^, but with a weaker level of conservation than some transcription factor binding sites in closer related Drosophila species^44^. Therefore, if a DNA element is an active HAS in one species but not in another species, it is likely that another active HAS is present nearby, such that the number of and spacing between HAS does not change drastically.

If the majority of HAS rapidly turn over throughout evolution, how do new HAS arise? We find that HAS are enriched in genic space, especially within introns and 3’UTRs (**Figure 5A** and **figure supplement 12A**). Enrichment on genic over intergenic regions is consistent with the idea that the DCC targets and regulates gene expression. Very few HAS are present in coding sequences, perhaps because the low complexity MRE motif is not well tolerated in ORFs^42^. As introns represent the most abundant location of roX binding (nearly half), we analyzed the position of HAS within introns. Notably, we found that HAS are proximal to the 3’-end of introns and are approximately 3-fold enriched at DNA encoding polypyrimidine tracts (PPT), a C/T-rich splicing signal (**Figure 5B** and **figure supplement 13A**). Approximately 20% of observed *D.melanogaster* HAS are within 100bp of a PPT, vs. ∼7% in a permuted background model (*p*-value=1.77e-11, K-S test). Conserved HAS are more enriched at PPT sites than species-specific HAS (28% vs. 15%). The association of HAS and PPT also holds true for *D.willistoni* and *D.virilis* (**figure supplement 12B-C**). Moreover, the 271 HAS on *D.willistoni* ME-D are significantly enriched near PPT (16% observed vs. 4% in permuted control, *p*-value<2.2e-16, **figure supplement 10**), despite ME-D’s ancestry as an autosome. Their homologous positions on the autosomal *D.melanogaster* ME-D are also enriched near PPT (29% observed vs. 7% in permuted control; *p*-value=3.90e-5, K-S test). Thus, after the fusion of an autosome and the X, HAS can evolve from ancestrally autosomal PPTs (**figure supplement 8**).

**Figure 5.**
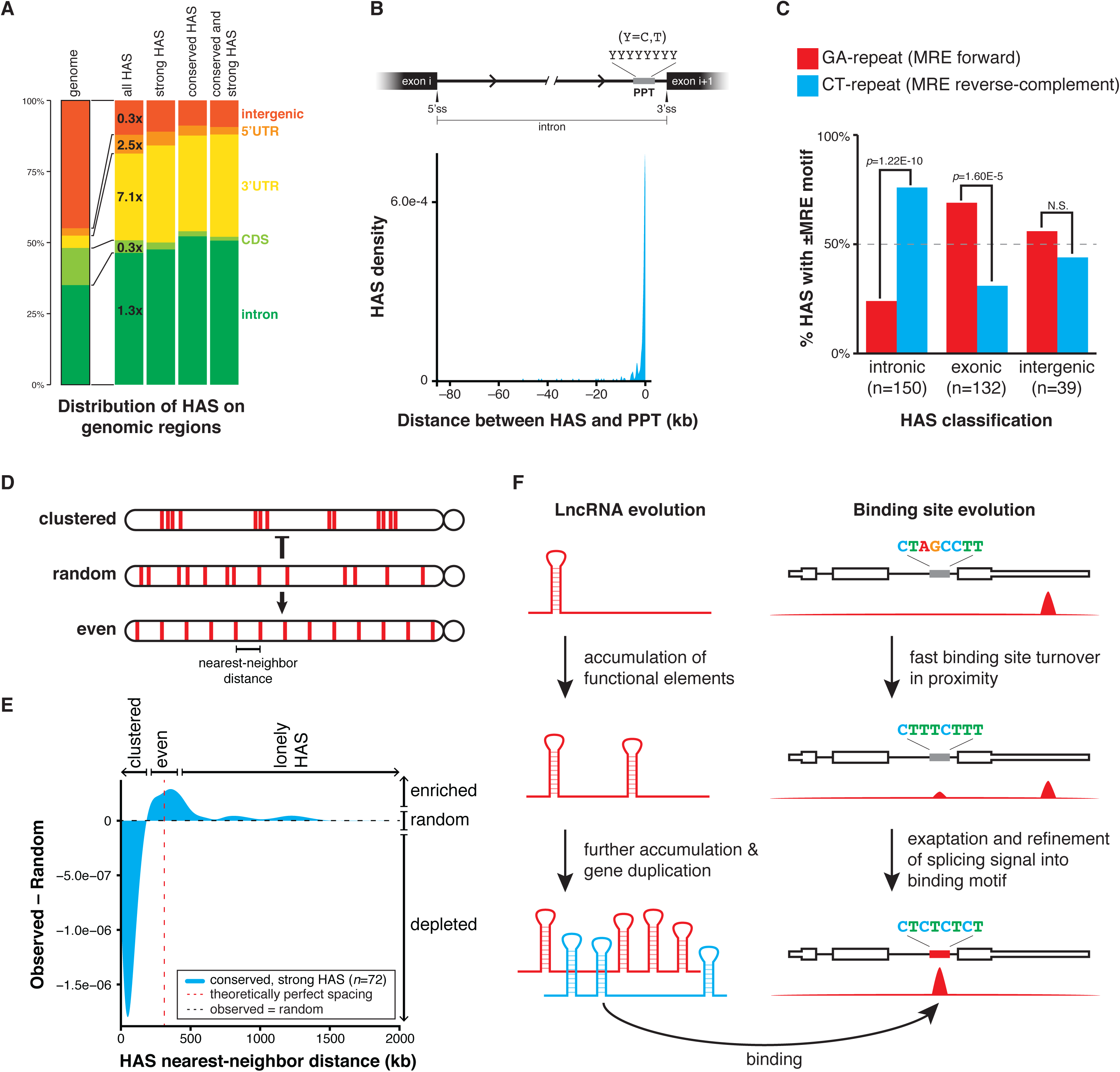
HAS exapt pre-existing regulatory signals and are selected for even spacing on the X-chromosome. **(A)** HAS are enriched in genic, noncoding regions of the genome, primarily within introns and 3’UTRs. HAS are subcategorized by strong roX occupancy and/or high evolutionary conservation. Fold enrichment over the genomic distribution is shown. See also **figure supplement 12A**. **(B)** Intronic HAS are proximal to polypyrimidine tracts (PPT). See also **figure supplement 12B-C**. **(C)** The MRE motif within HAS classes exhibit significant orientation bias. Intronic HAS are biased in the reverse-complement orientation (CT-repeat), whereas exonic HAS are biased in the forward orientation (GA-repeat); intergenic HAS have no bias. See also **figure supplement 12D**. **(D)** Alternative HAS spacing models on the X-chromosome. HAS may be clustered together, randomly spaced, or evenly spaced. The observed distribution is a mix of randomly and evenly spaced distributions. **(E)** The difference between the observed and random HAS (conserved, strong only) distributions on the X-chromosome. The positive *y*-value near the theoretically perfect spacing distance indicates an enrichment of the even spacing model relative to random spacing; conversely, the negative y-value at short distances indicates a depletion of the clustered spacing model relative to random spacing. (**F, left**) lncRNAs like roX can evolve function through the accumulation of repetitive structural or sequence elements. (**F, right**) HAS are evolutionarily dynamic, losing function at one genetic element while gaining function nearby, and can co-opt existing polypyrimidine tracts within gene introns as a source of MRE elements.

The reverse-complement of the GA-repeat MRE motif (CT-repeat) closely resembles the C/T-rich sequence of PPT, raising the hypothesis that PPT may serve as an abundant evolutionary source of MRE-precursors. To test this hypothesis we measured the strand bias in the MRE motif orientation relative to the direction of gene transcription. In the null hypothesis, MRE motifs in DNA would be independent of transcriptional direction and have no strand bias. Conversely, the PPT hypothesis predicts that MRE motifs would be biased towards the pyrimidine-rich orientation. Indeed, intronic HAS are significantly over-represented by the reverse-complement MRE motif (CT-dinucleotide repeat; *p*-value=1.22E-10, binomial test; **Figure 5C** and **figure supplement 12D**), and HAS-containing introns are more pyrimidine-rich (*p*-value=4.21e-5, K-S test; **figure supplement 13B**) and shorter than typical introns (*p*-value<2.2e-16, K-S test; **figure supplement 13C**). Taken together, these results suggest that some PPT motifs moonlight as MRE, coopted for dosage compensation and evolutionarily refined into HAS (**Figure 5E** and **figure supplement 14**). Curiously, we also found that exonic HAS (primarily in 3’UTRs) are significantly over-represented by the forward MRE motif (GA-dinucleotide repeat; *p-*value=1.60E-5, binomial test; **Figure 5C** and **figure supplement 12D**). This bias further distinguishes PPT from other transcriptional units and reflects the slightly purine-rich environment of exons (**figure supplement 13D-E**).

Finally, we addressed potential selective pressures that drive the conservation of a subclass of HAS. We did not find any obvious genomic features or gene ontology terms for the genes near HAS with the highest evolutionary conservation and strongest binding signal. However, these “conserved, strong” HAS (72 in *D.melanogaster*) are more evenly spaced along the X-chromosome than expected by chance alone (**Figure 5D-E** and **figure supplement 12E**). The distribution of distances between nearest-neighbor HAS is different from permutation tests with the same number of HAS, and is enriched near the theoretically perfectly spacing distance (the length of the X-chromosome divided by the number of HAS). The more-even-than-random placement of HAS thus maximizes HAS distribution along the X, which may therefore allow the DCC to spread as effectively as possible from a minimal number of HAS. As the dosage compensation complex also participates in organizing higher-order chromosomal looping^37, 45^, we speculate that chromosome architecture or other roles may drive the selection for a subset of HAS at evenly spaced locations.

## Discussion

Using an integrative “nested homology” strategy based on phylogenetic conservation of synteny, microhomology, and RNA structure, we successfully identified 47 previously unknown roX lncRNA orthologs from 35 diverse flies. Despite very poor primary sequence homology, these distantly related roX orthologs have conserved function and can suffice for dosage compensation in *D.melanogaster*. The discovery of these diverse roX orthologs permitted comparative analyses of RNA sequence, structure, and genomic interactions, revealing principles of lncRNA evolution and genomic targeting (**Figure 5F**). This integrative approach is likely applicable to trace the evolutionary dynamics of many lncRNAs that populate all kingdoms of life, as demonstrated by our description of the *HOTAIR* locus in species as diverse as human and zebrafish (**figure supplement 15**). Furthermore, recent methods that reveal RNA structures *in vivo*^46^ should facilitate the systematic organization of lncRNAs by structural homologies. The search strategy described here differs from others in that it is targeted in scope and only requires query genomes, whereas others are dependent on RNA-seq datasets, which are often sparse for non-model organisms^11, 12^.

Focal structures and repeated sequences emerged as key features for both the discovery and function of roX lncRNAs. In distantly related species, the roXbox stem-loops are often the only recognizable features linking roX RNA orthologs, and the number of the repeats correlates with the ability of roX1 orthologs to occupy the X-chromosome. This insight also allowed us to engineer designer lncRNA transgenes with one or more roXbox stem-loops, which functioned to varying degrees *in vivo* (**Figure 3C-D**). This fits with the concept that lncRNAs evolve rapidly and can act as flexible scaffolds tethering together one or more functional elements. We found evidence for roX gene duplication in some species, producing lncRNA paralogs with support for partial lncRNA “pseudogenization” of one paralog (**figure supplement 1**). Similarly, we showed that the complete roX1–roX2 functional redundancy observed in *D.melanogaster* is likely unique to certain species within the Sophophora subgenus, as roX1 orthologs in the Drosophila subgenus have lower expression, limited localization to the X-chromosome, and systematic loss of key structures and domains. The function, if any, of roX1 in the Drosophila subgenus may be addressed in the future by genetic disruption of one or both *roX* genes. Additionally, the discovery of roX1 and roX2 orthologs in more distantly related outgroup species may shed light on the evolutionary origin of these lncRNAs, but would require more fully sequenced fly genomes. Similarly, did roX1 and roX2 originally evolve from an ancestral roX gene duplication event? Perhaps roX1–roX2 functional redundancy in certain flies allows divergent specialization in their regulatory programs or expression patterns, as with duplicated protein-coding genes that acquire divergent roles^47^. The repetition and refinement of functional elements may be a general principle in the evolution of some lncRNAs, as with roX and XIST (**Figure 5F**). Tracing the evolutionary patterns of key sequence or structural elements may shed light on the origin, diversification, and extinction of lncRNA genes.

Genome-wide roX occupancy maps in several species revealed the evolutionary constraints on lncRNA–genome interactions. roX binding sites are always strongly enriched on the X-chromosome, can turn over quickly, and are constrained in their local chromosomal neighborhood and spacing pattern (**Figure 5F**). This pattern of evolutionary conservation is reminiscent of enhancer elements that bind transcription factors^44^. The even spacing pattern of binding sites maximizes spreading while simultaneously minimizing the total number of HAS. Moreover, prior studies in *D.melanogaster* suggested that roX can spread by spatial proximity in 3D rather than linearly^37, 45^, which is consistent with the conservation of roX binding sites in local spatial neighborhoods. Our discovery of rapid turnover of individual roX binding sites implies that new HAS must be born frequently, such that mutations of existing HAS do not compromise X-chromosome dosage compensation and can invade neo-X chromosomes rapidly. Furthermore, the evolutionary dynamism of HAS implies that DCC action is distributed rather than targeted, with the primary constraint being that targeting is to many interspersed sites on the X-chromosome, but not necessarily specific genes or sites on genes. One abundant source of new HAS are intronic PPT (**Figure 5F** and **figure supplement 14**), a feature of lncRNA targeting that was not previously appreciated, which would further facilitate the rapid invasion of the DCC to neo-X chromosomes. Exaptation of this splicing signal is an elegant strategy for dosage compensation because it parsimoniously encodes one function at the level of DNA (DCC-binding) and another at the level of RNA (splicing). Additionally, coupling nascent roX targeting to DNA sequences encoding an RNA splicing signal may ensure that the dosage compensation machinery is targeted to bona fide genes that are actively transcribed and spliced. Comparative genomic studies of lncRNAs and their binding sites will be a powerful approach to address these and other questions about the noncoding genome in the future.

**Supplementary Information** is available in the online version of the paper.

## Acknowledgements

We thank members of the Chang and Akhtar labs for discussion. Supported by NIH grants and Howard Hughes Medical Institute (H.Y.C.), Max Planck Society (A.A.), and Bio-X Fellowship (J.J.Q.).

## Author Contributions

J.J.Q., Q.C.Z., and H.Y.C. conceived the project. J.J.Q. carried out experiments and lncRNA search; I.A.I. and P.G. performed genetic experiments. Q.C.Z. and J.J.Q. performed computational analyses. J.J.Q. and H.Y.C. wrote the paper with input from all authors.

## Author Information

The authors declare no competing financial interests. All raw and processed sequencing data can be accessed in NCBI’s Gene Expression Omnibus through accession number GSE69208.

## Materials & Methods

### lncRNA ortholog search strategy

The general principle for the lncRNA search strategy follows three primary steps: (1) search initiation with a known lncRNA, (2) searching for closest-relative lncRNA orthologs using synteny, microhomology, and/or structure features, and (3) iteratively refining the search parameters with each newly discovered lncRNA ortholog and searching for the next-closest-relative lncRNA ortholog. In this way, to complete this search, one only needs knowledge of an initiating lncRNA from a single subject species (features such as its sequence and neighboring genes), sequenced genomes of other query species, and the phylogenetic relationships between the subject species and the query species. In some instances, a sequenced genome is not necessary for discovering new lncRNA orthologs, as described below (i.e. analog search strategy based on degenerate PCR of syntenic protein-coding genes).

To initialize the search, we collected knowledge of roX1 and roX2 in *D.melanogaster* (and HOTAIR in *H.sapiens*), specifically the neighboring syntenic genes, instances of repeated microhomology, and known secondary structures – both measured and predicted^21, 37^. For example, in *D.melanogaster*, *roX1* is flanked by protein-coding genes *yin* (upstream, sense) and *ec* (downstream, antisense); *roX2* is flanked by protein-coding genes *e(y)2* (upstream, antisense), *CG11695* (upstream, sense), and *nod* (downstream, sense) (**figure supplement 2**). Human HOTAIR is encoded in a ∼17kb window between protein-coding genes *HOXC11* and *HOXC12* (**figure supplement 15**). To find repeated microhomologous sequence elements shared between roX1 and roX2, we searched for matching motifs shared between both RNAs using MEME^50^ (*site distribution*: any number of repetitions); this returned a sequence motif collapsing roXboxes from roX1 and roX2. The structures of roX1 and roX2 have been measured or predicted previously^21, 37^, and we used NUPACK-predicted^51^ structures for visual comparison to other lncRNA ortholog candidate structures.

Using tBLASTn, we used the amino acid sequences of syntenic *D.melanogaster* protein-coding genes in D.melanogaster to search for orthologous protein-coding genes in the closest-relative fly species (e.g. *D.simulans, D.sechellia, D.mauritiana, D.yakuba,* and *D.erecta*). This then defined the genomic interval surrounding the candidate orthologous *roX* loci. Next, using the collapsed roXbox sequence motif from *D.melanogaster* roX1 and roX2, (as a position-weight matrix from MEME^50^), we matched the motif to sites within the synteny block (using FIMO^50^). In each case, this elected a ∼500bp window with a cluster of 3-6 high-scoring roXbox incidences, corresponding to roX1-D3 domain or roX2 exon-3^21, 37^. We also computed the minimum energy structures within these windows (using NUPACK^51^), and visually compared the predictions to the structures in *D.melanogaster* roX1 and roX2, such as the repeated roXbox stem-loops^21, 31, 37^.

Using these new high-confidence roX1 and roX2 ortholog candidates from the expanded species list (i.e. all melanogaster subgroup flies), we next refined the search parameters. Neighboring syntenic genes remained unchanged, but we updated the microhomologous motifs with the additional roX1 or roX2 orthologs (thus improving the accuracy of the motifs and finding additional weakly conserved sites that could also be used for the orthology search). The minimum free energy structures for each of these species’ RNAs were collated for comparison in the next iterative rounds of the search strategy. Equipped with these refined search parameters, we expanded the search to more distantly related flies, such as those in the melanogaster group, thus iterating the search strategy and leveraging known phylogenetic relationships. For example, though *roX2* neighbors the *nod* gene in *D.melanogaster*, *roX2* neighbors *ari-1* in flies outside of the melanogaster subgroup (e.g. *D.takahashii*, **figure supplement 2**); thus, we abandoned searching for syntenic regions around *nod* in flies outside of the melanogaster subgroup and instead used *ari-1*. With each new lncRNA ortholog candidate discovered, the search parameters become more and more refined, thus enabling the discovery and more distantly related orthologs.

In species lacking WGS assemblies, we used a PCR-based method to perform the synteny search. We designed degenerate PCR primers at conserved sequences in protein-coding genes expected to be syntenic with *roX* RNAs; if synteny was preserved, PCR yielded a DNA fragment, which we sequenced and then proceeded with the search strategy. By syntenic PCR, we found roX1 in *D.nasuta*, but not *D.guttifera, C.pararufithorax,* nor *C.amoena;* this suggests that either *ec*–*yin* synteny blocks have been disrupted or the syntenic protein-coding gene sequences are too divergent. We did not search for roX1 in *D.paulistorum, D.nebulosa, D.saltans*, or *H.duncani*, as these flies were included for studying roX2–roX3 paralogy and lack WGS. roX2 could not be identified in *Scaptodrosophila lebanonensis* because the *e(y)2–ari-1* loci are incompletely scaffolded due to low N50 of this genome assembly.

### Fly species and rearing

All fly stock species were sourced from the Drosophila Species Stock Center (stockcenter.ucsd.edu); the species stocks used here are listed in the **Supplementary Information**. All flies were raised on standard cornmeal-molasses medium or Wheeler-Clayton medium (*D.busckii* only) at room temperature, unless specified otherwise.

For genetic experiments, the following stocks were obtained from the Bloomington stock center or were kindly donated: *y*^*1*^ *w*; P{tubP*-*GAL4}LL7/TM3, Sb1* (Bloomington stock #5138), *w*^*1118*^;*P{da*-*GAL4.w*^-^*}3* (Bloomington stock #8641), *roX1^SMC17A^, roX^2Δ^; CyO, hsp83*-*roX1* (ref. ^52^).

### Genomic DNA and crude RNA extraction

Genomic DNA was extracted from whole adult mixed-sex flies using the Gentra Puregene kit (Qiagen); the gDNA was used for validation of roX loci sequences from WGS or synteny PCR with degenerate primers, as listed in the **Supplementary Information**. Crude RNA was extracted from whole, newly-eclosed male or female flies using TRIzol reagent (Life Technologies), treated with TURBO DNase (Life Technologies), and cleaned-up on RNeasy Mini columns (Qiagen); the crude RNA was used for RT-PCR expression analysis and RACE.

### Polytene squashes and RNA FISH

Polytene chromosome squashes were prepared from sexed wandering third instar larvae. First, larvae were washed gently in PBS; under a dissection microscope, larvae were inverted and the salivary glands were dissected while carefully removing the attached fat bodies. The salivary glands were then fixed first in a depression slide with 100μL of 3.7% formaldehyde + 1% Triton X-100 in PBS for 45 seconds, followed by 3.7% formaldehyde in 50% acetic acid for 2 minutes. The salivary glands were then transferred to 15μL of a 50% acetic acid, 17% lactic acid solution on a siliconized coverslip and immediately inverted onto a polylysine microscope slide. The polytene chromosomes were spread and squashed by gentle tapping and wiggling of the coverslip and checked under a phase microscope. Excess solution was removed with a clean wipe. The slide was then flash frozen in liquid nitrogen, the coverslip was removed, and the polytene chromosomes on the microscope slide were dehydrated in 100% ethanol for 30 minutes. Finally, the slides were washed twice in PBS before proceeding to single molecule FISH staining and imaging on a fluorescent microscope, according to the Stellaris protocol (Biosearch Technologies). Single molecule FISH probes are listed in the **Supplementary Information**.

### RT-PCR, RACE, and synteny PCR

Oligo(dT)-primed cDNA libraries were made from crude RNA extract from each species using SuperScript III First-Strand Synthesis System (Life Technologies). RT-PCR was performed using species-specific primers against roX1, roX2 (and roX3, where possible), and GPDH, and amplified for a total of 30 cycles from 1μL of the starting cDNA library. 5’- and 3’-RACE were performed using the GeneRacer kit (Life Technologies) starting from crude RNA. Syntenic PCR was performed from genomic DNA and using degenerate primers designed against conserved syntenic genes or regions. See **Supplementary Information** for the lists of all primers used.

### Sequence identity and structure modeling

Sequence conservation was calculated using Clustal Omega 1.2.1 (DNA, standard settings) for each roX1, roX2 (and roX3, where applicable), and roX2 intron relative to two independently scrambled sequences, generated by scrambling the *D.melanogaster* sequence. The pairwise % sequence identity was calculated for every pairwise comparison. The lower limit of sequence homology was determined by averaging the pairwise % sequence identity between each gene and the scrambled sequences, and was ∼36% (not the theoretical 25%, since nucleotides are not evenly represented in the roX RNAs). Pairwise % sequence identity was then graphed using the upper and lower bounds of 100 and the scrambled percentage. NUPACK^51^ was used to predict local RNA secondary structures in roX1 and roX2, modeled after the experimentally validated structures in *D.melanogaster*^21^.

### Genetic Experiments

Fly work has been done essentially as described in Ilik et al., 2013 and Quinn et al., 2014. Briefly, all *roX1* and *roX2* constructs were cloned into pUASattB vector and transgenic flies were generated using phiC31 integrase-mediated germ-line transformation as previously described^53^, injecting *y*^*1*^ *M{vas*-*int.Dm}ZH*-*2A w*; PBac{y^+^attP- 3B}VK00033* embryos. To score male-specific lethality rescue, *roX1SMC^17^A, roX^2^^Δ^ daGAL4* or *roX1^SMC17A^, roX^2^^Δ^ tubGal4/TM6Tb* virgin females were crossed to *UAS-roX1\** and *UAS-roX2\** males, respectively, and allowed to develop at 25°C. *roX1*\* denotes the transgenic construct, namely *D.mel roX1* (full-length), *D.vir roX1-D3*, and *D.vir roX1-D3 +D.mel SL* in **Figure 3D**. *roX2*\* denotes the transgenic construct, namely *D.mel roX2-exon3*, *D.bus roX2-exon3*, *D.bus roX2-5’ + D.mel roX2-3’* (Chimera 1), and *D.mel roX2-5’ + D.bus roX2-3’* (Chimera 2) in **Figure 2E**.

Male and female adult flies from at least three independent crosses were counted daily for a period of 10 days from the start of eclosion, without blinding. The total number of non-Tb males was divided by the total number of non-Tb females that eclosed during the 10-day period, which was used as an internal control for 100% viability.

Gene expression analysis was done as described previously^21, 37^. Briefly, 3-4 3^rd^ instar larvae were homogenized in Trizol and total RNA was extracted using the Direct-zol kit (Zymo). RNA was reverse transcribed with SuperScript III and random hexamers (Life Technologies). Relative expression values were calculated using the 2ΔΔ^Ct^ method, using phosphofructokinase RNA as the internal control.

### *In vivo* ChIRP-seq

ChIRP-seq protocol was adapted from Chu et al., 2011 and chromatin preparation from larvae was adapted from Soruco et al., 2013 and Alekseyenko et al., 2006. First, 1.0 gram of mixed-sex wandering third instar larvae (between ∼300-1500 larvae, depending on the species) were collected, washed in PBS, flash-frozen in liquid nitrogen, and pulverized into a fine powder using a mortar and pestle under liquid nitrogen. Next, the powder was reconstituted in 40mL cold PBS with protease inhibitor cocktail (Roche) and homogenized in a dounce tissue grinder (Kimble Chase) with 5 passes of douncer A and 2 passes of douncer B. The homogenized material was passed through a 100μm nylon SteriFlip filter (Millipore) and immediately fixed in 1% formaldehyde (Thermo Scientific) by nutation for 20min at room temperature. Fixation was then quenched by adding 5% volume of 2.5M glycine and nutating for 5min at room temperature. The fixed material was pelleted by centrifugation at 3800RPM for 30min at 4°C, and the pellet was washed with cold PBS and pelleted. The pelleted material was resuspended in 2mL cold Swelling Buffer (0.1M Tris pH 7.0, 10mM KOAc, 15mM MgOAc) supplemented with 1% NP-40, protease inhibitor, PMSF, and Superase-In (Ambion), incubated on ice for 10min, and dounced for 2sec with a hand-held motorized homogenizer (Argos) fitted with 1.5mL tube pestles (VWR). Material was pelleted by centrifugation at 5000rcf for 10min at 4°C and washed in cold PBS with 0.5mM EDTA. Next, the material was further fixed with 3% formaldehyde in PBS by nutation for 30min at room temperature; cross-linked material was pellete by centrifugation at 3500RPM for 30min at 4°C, washed in PBS, and pelleted. Cross-linked material was resuspended in 7mL Nuclear Lysis Buffer supplemented with protease inhibitor cocktail, PMSF, and Superase-In, and then solubilized and sheared by sonication using a Covaris E-series focused ultrasonicator (850μL per tube; 4°C water bath; 5% duty cycle; 140 PIP; 60min total sonication time). Nucleic acid shearing was confirmed by agarose gel electrophoresis. The resulting chromatin was clarified by centrifuging at maximum speed on a tabletop minifuge for 10min at 4°C; the soluble chromatin fraction was collected without disturbing the insoluble pellet, and flash-frozen in liquid nitrogen or immediately used for ChIRP.

ChIRP was performed as described in Chu et al., 2011. ChIRP oligos were designed against roX1 and roX2 RNAs from each species using the Stellaris single molecule FISH oligo designer (Biosearch Technologies), as previously described^37^. ChIRP oligos are listed in the **Supplementary Information**. The DNA fraction from each ChIRP experiment and inputs were purified and libraries were constructed using the NEBNext DNA Library Prep kit (NEB). Sequencing libraries were barcoded using TruSeq adapters and sequenced on HiSeq or NextSeq instruments (Illumina) using single-end reads of 50bp length (1x50). Reads were processed using the ChIRP-seq pipeline as previously described^24^.

### Peak calling, filtering, and motif analyses

Peaks were called from the merged [=Minimum(Even,Odd)] roX2 ChIRP-seq tracks using MACS2 (no peak model; 150bp extension size; summit calling enabled). Called peaks were filtered by their significance (- log_10_(*q*-score) ≥3000; ≥8000 for *D.willistoni*) and enrichment (ChIRP/input ≥20).

Sequence motifs were discovered from the 500bp windows centered around peak summits using MEME (zero or one occurrence per sequence; 21bp window). The central location of each motif occurrence was determined using CentriMo^50^.

### Signal enrichment analysis

ChIRP-seq signal enrichment was calculated for every 1kb window of the genome as the sum of signal from roX1 or roX2 ChIRP divided by the input signal from the same window. The enrichment was then plotted as grouped by Müller element assignments (see below). The 5kb windows around the *roX1* and *roX2* loci were excluded, due to the possibility of direct genomic DNA recovery by antisense ChIRP oligos. To calculate the roX1 vs. roX2 signal bias, the ratio of roX1 to roX2 ChIRP-seq signal was calculated for all called peaks. Box-and-whisker plots represent the 95, 75, 50, 25, and 5^th^ percentiles, plotted on a log_2_ scale, and the fractional bias represents the median roX1-to-roX2 bias.

### Genome assemblies

All genome builds were obtained from the Flybase (www.flybase.org), with exceptions. The genome assembly for *D.americana* was downloaded from the Jorge Vieira lab website (evolution.ibmc.up.pt); *D.suzukii* from the Spotted Wing Flybase (spottedwingflybase.oregonstate.edu; ref. ^54^); *D.mauritiana* from popoolation.at/mauritiana_genome/index.html (ref. ^55^). For *D.buskii*, we downloaded the raw WGS reads from the NCBI Short Read Archive (SRP021047), which contains 90bp paired-end reads from female flies as described^38^. We trimmed the data, and assembled the genome as described with the exception of using SOAPdenovo2^56^, the updated version of SOAPdenovo. Only scaffolds longer than 1kb were kept for further analysis. Our assembly of the *D.busckii* genome is available from GEO; see accession below. The genome assembly statistics are in **Supplementary Information**.

### Protein-coding gene annotation

We obtained all genome annotations from Flybase (www.flybase.org), except for *D.buskii*. The genome annotation information is available from GEO; see accession below. For *D.buskii*, we annotated its putative protein-coding genes by using homology transfer of *D.melanogaster* protein-coding sequences, downloaded from Flybase. The homology transfer was based on the genBlastA pipeline^57^, which uses BLAST to find high-scoring pairs (HSPs) between *D. melanogaster* and *D.buskii*, and then uses a graph-based algorithm to filter and select an HSP group representing a homologous gene of *D. melanogaster* in *D.buskii*. The parameters used in running genBlastA are: -p T -e 1e-1 -g T -f F -a 0.6 -c 0.4 -d 100000 -r 10 -s 0.

For each ChIRP-seq peak, we then used the tool *intersectBed* in the *BEDTools* suite^58^ to find the genomic feature(s) to which the peak summit belongs, based on Flybase annotations. A small fraction of genomic features overlap, and as such some peak summits were counted separately. For example, a peak summit could be in the intron of one transcript and the exon of another, and thus the peak will be counted twice.

### Müller element annotation

We implemented a pairwise genome alignment pipeline based on LASTZ^59^, and the UCSC tool set. We followed a description from the UCSC website (genomewiki.cse.ucsc.edu/index.php/Whole_genome_alignment_howto), and used the same parameter settings by UCSC (hgdownload.soe.ucsc.edu/goldenPath/dm3/vsDroVir3/). We compared our alignment of *D.melanogaster* and *D.virilis* with the liftOver file downloaded from UCSC and confirmed that they are virtually identical.

Based on the pairwise genome alignment, we were able to calculate an empirical similarity score for each scaffold of *D.virilis*, *D.willistoni*, and *D.buskii* and each Müller element of *D.melanogaster*. The score is defined as the chain score between the scaffold in the first species and the Müller element in the latter divided by the total chain score of the scaffold and all Müller elements. We applied a stringent cutoff of 0.85 to reliably assign a scaffold to a Müller element. This assigns most scaffolds that are longer than 1000. For the very long scaffolds that are not assigned by this cutoff, we manually inspected the empirical score and also the homology information of protein-coding genes on the scaffolds and the correspondent Müller element. For example, we assigned *D.willistoni* scaffold scf2_1100000004963 to Müller element A (similarity score: 0.797, protein homology percentage: 90%, i.e. 90% proteins of scf2_1100000004963 are homologous proteins in *D.melanogaster* Müller element A).

### Gene-level and Element-level peak overlaps

For each ChIRP-seq peak, we assigned a gene association if the peak summit was within 1kb of the gene. For *D.virilis*, *D.willistoni*, and *D.buskii*, since the UTR regions were not usually annotated, we included a typical length of 200bp or 500bp for the 5’UTR and 3’UTR, respectively. After this assignment, starting from each peak in each species, we asked whether it had related peaks in other species, based on the orthology information annotated in Flybase for *D.melanogaster* genes and *D.virilis* or *D.willistoni* genes, and from our own genBlastA pipeline annotation for *D.melanogaster* and *D.busckii* (described above). For a peak in species *A*, if its associated genes contained a gene ortholog associated with a peak in species *B*, the peak was regarded as gene-wise conserved between species *A* and *B*. A peak was called gene-wise invariant if it was conserved among all four species. On the contrary, if no associated gene of a peak in species *A* was an ortholog of any associated gene in any other species, the peak was regarded as species-specific to species *A*.

We also investigated the conservation of the genomic positions of a ChIRP-seq peak in different species based on our pair-wise whole genome alignment. Specifically, for each peak in a species *A*, we used the liftOver tool to find its homologous position in species *B*. If the position overlapped with a peak in species *B*, it was regarded as conserved. We studied the peak turnover by allowing the homologous position to be within certain distance of a peak in species *B*. We observed that if the homologous position did not overlap with a peak in species *B*, there was often a peak present nearby. We compared this distribution to random chance by permuting the peaks on species *B* within the same chromosome or across the whole genome, by using the *shuffleBed* tool from the *BEDTools* suite.

### Peak-to-PPT summit calculation

For each intron we obtained its sequence and predicted the positions of polypyrimidine tracts (PPTs) within the intron by using the online tool SVM-BPfinder^60^. We then implemented an algorithm to select the most likely PPT for each intron, by adding a penalty score, which increases with the distance to the 3’ splicing site, to the final score of SVM-BPfinder. Specifically, if the distance was <40bp of the 3’ splicing site, the penalty score equals 0, but increase 0.02 for each base in distance. For all ChIRP-seq peaks, we calculated the directional distance to its nearest PPT (“–“ means that the peak summit was upstream of the intron’s PPT, and “+” means downstream).

We then permuted the position of each ChIRP-seq peak within the same chromosome and calculated again the directional distance of a random peak to its nearest PPT. We compared the two distributions by using a two-tailed K-S test. We also counted the number and percentage of observed or random peaks within 100bp of a PPT.

### MRE motif orientation bias analysis

We used MEME^50^ to identify the position and orientation of the best MRE motif within each ChIRP-seq peak of each species. The positions of the MRE motif were used to annotate which genomic feature the peaks were then assigned (e.g. CDS, intron, etc.). The motif orientation instances were counted for each category of genomic features, and a binomial test was used to quantify the differences.

### Chromosome spacing analysis

We calculated the distance for each peak summit to its nearest neighbor. If ChIRP-seq peaks were perfectly evenly distributed on a chromosome, the nearest-neighbor distances would always be the length of the chromosome divided by the total number of peaks. If all peaks were clustered into a few hot spots, the nearest-neighbor distances will approach zero. We also simulated the random distance distributions by shuffling the peaks to random positions within the chromosome.

We defined a subset of strong peaks (enrichment>50 or log_10_(q-value)>10000; >20000 for *D.virilis*) or conserved peaks (if a peak overlapped another peak in a second species). We further defined a subset of strong and conserved peaks as the intersection of these two sets. We calculated the above analysis of nearest-neighbor distance using this subset of peaks. Plotted is the difference between observed and the random distributions of nearest-neighbor peak distances.

### Supplementary Information

The accompanying **Supplementary Information** contains the list of RT-PCR primer sequences, 5’- and 3’-RACE primer sequences, degenerate synteny PCR primer sequences, ChIRP oligo sequences, qRT-PCR primer sequences, fly species stock numbers, and the genome assembly releases used.

### Accession codes

The raw sequencing reads from each ChIRP-seq experiment (*.fastq), the mapped and merged ChIRP-seq and input tracks (*.bedGraph and .bigWig), called ChIRP-seq peaks (*.bed), Müller element assignments (*.xlsx), *D.busckii* genome assembly and annotations, and roX1 and roX2 sequences can be accessed at GEO (accession number: GSE69208).

**Figure Supplement 1.**
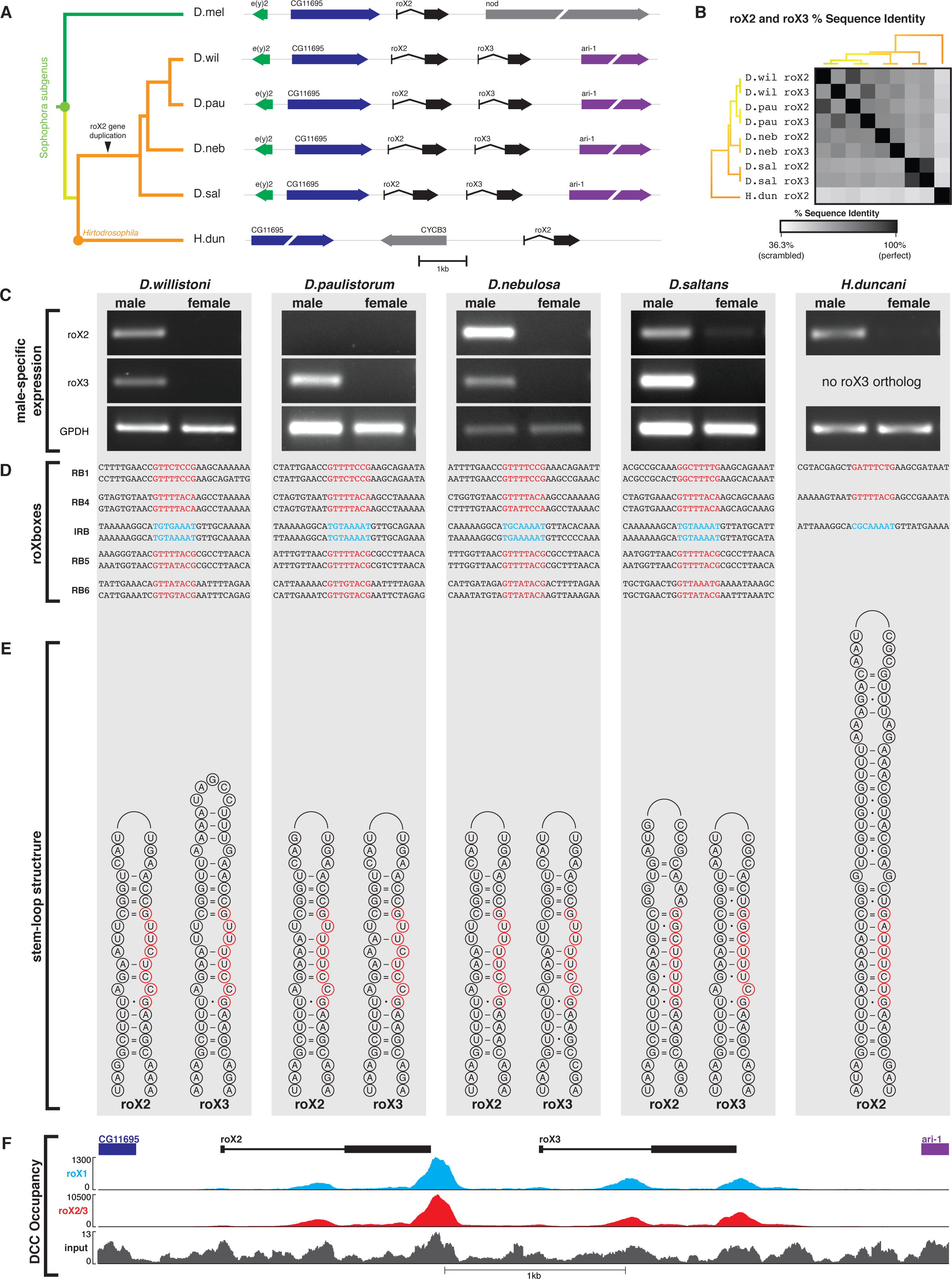
*D.willistoni* has three roX RNAs resulting from a *roX2* gene duplication event in the *willistoni-saltans* clade. (A) The *roX2–roX3* locus of *D.willistoni* and its relatives. Our ortholog search in *D.willistoni* identified a third roX homolog candidate adjacent to *roX2*. Using synteny PCR from *e(y)2* and *ari-1* or *roX2*, we cloned and sequenced the *roX2* – *roX3* locus in four relatives of *D.willistoni*. The roX2–roX3 pairs are present in *D.willistoni, D.paulistorum, D.nebulosa*, and *D.saltans*, but absent in the sister species *Hirtodrosophila duncani*, indicating that roX3 arose after the divergence of *H.duncani* and the concestor of the *willistoni*-*D.saltans* clade. The *D.melanogaster roX2* locus is shown for reference. (B) Sequence identity between roX2 and roX3 in five species. The roX2 and roX3 orthologs share relatively high sequence identity both between species and between roX2–roX3 pairs, indicating that roX2 and roX3 are likely paralogs that resulted from a whole gene duplication event. (C) roX2 and roX3 RNA expression. roX2 and roX3 orthologs are male-biased transcripts in the *willistoni-saltans* clade, as is roX2 in *H.duncani*. Expression bias between roX2 and roX3 is species-specific; for example, roX2 expression is nearly undetectable in *D.paulistorum*, yet roX2 expression is higher than roX3 in *D.nebulosa*. GPDH is used as a sex-independent control. (D) The roXboxes (RB1, RB4-6; red) and inverted roXbox (IRB; cyan) of roX2–roX3 orthologs are conserved horizontally between species and vertically between roX2–roX3 paralogs. (E) The stem-loop structure at the 5’-end of roX2 and roX3 orthologs is conserved between species and between roX2–roX3 paralogs. This structure contains RB1 (red circles). *H.duncani* roX2 has an extended stem-loop. (F) ChIRP-seq was used to map the chromatin occupancy of the roX RNAs in *D.willistoni*. The *roX2–roX3* locus is shown with signal from roX1 and roX2/3 ChIRP-seq and input DNA. The *roX2* and *roX3* loci exhibit a very similar pattern of roX binding, indicating that the HAS at these paralogous loci are also conserved.

**Figure Supplement 2.**
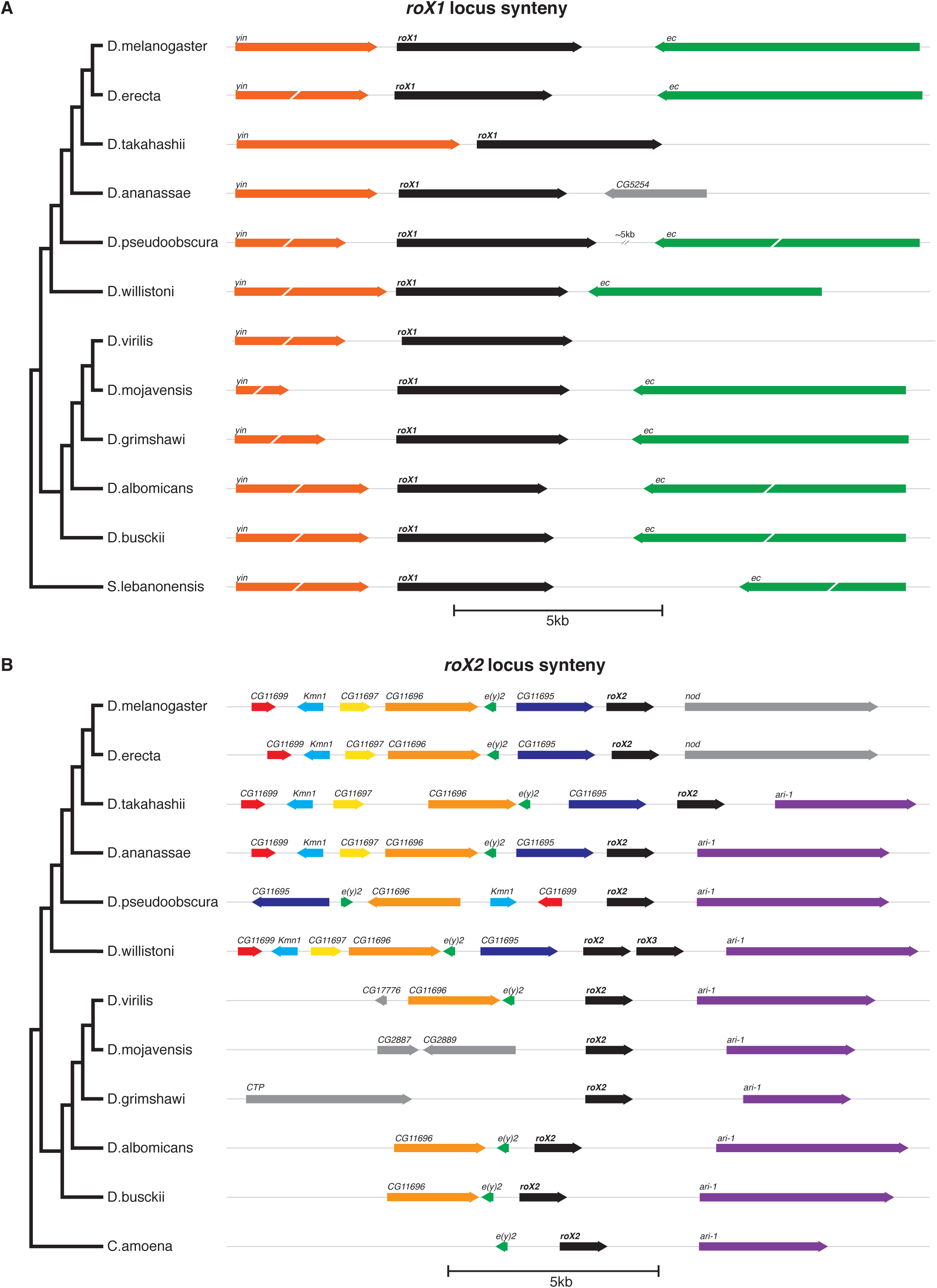
roX1 and roX2 ortholog synteny is maintained across evolution. (A) In *D.melanogaster*, *roX1* is flanked by protein-coding genes *yin* (upstream, sense) and *ec* (downstream, antisense). In nearly all other species, these flanking genes are maintained in position and orientation; one such exception is *D.ananassae*, in which an intrachromosomal rearrangement replaced the downstream neighboring gene *ec* with *CG5254*. For brevity, only representative species from major fly clades are shown. Scale bars, 5kb. (B) Similarly, in *D.melanogaster*, *roX2* is flanked by protein-coding genes *CG11695* (upstream, sense), *e(y)2* (upstream, antisense), and *nod* (downstream, sense). Synteny is not always perfectly preserved, however; though *nod* is downstream of *roX2* in the *melanogaster* subgroup, *ari-1* is downstream in all other flies outside the *melanogaster* subgroup. Thus, the *roX2*–*ari-1* synteny block is ancestral, and an intrachromosomal shuffling event in the ancestor of the *melanogaster* subgroup replaced *ari-1* with *nod* at the *roX2* locus. For this reason, when searching for roX2 orthologs in species outside the *melanogaster* subgroup, we instead used *ari-1* as a flanking gene instead of *nod*. Using *nod* in any search outside of the *melanogaster* subgroup would prove unproductive, since this syntenic relationship is not maintained. This instance highlights the importance of using phylogenetic relationships as a guide in the search strategy. When performing the synteny search in species lacking WGS, such as *D.nasuta*, we anticipated that *D.nasuta* might have similar gene order to its closest relative with a sequenced genome, *D.albomicans*. We designed degenerate primers at evolutionarily conserved sites in the four flanking protein-coding genes (*ec* and *yin*, and *e(y)2* and *ari-1*, for the *roX1* and *roX2* loci, respectively). In both cases, PCR yielded a fragment of roughly the expected intervening size, which we then sequenced.

**Figure Supplement 3.**
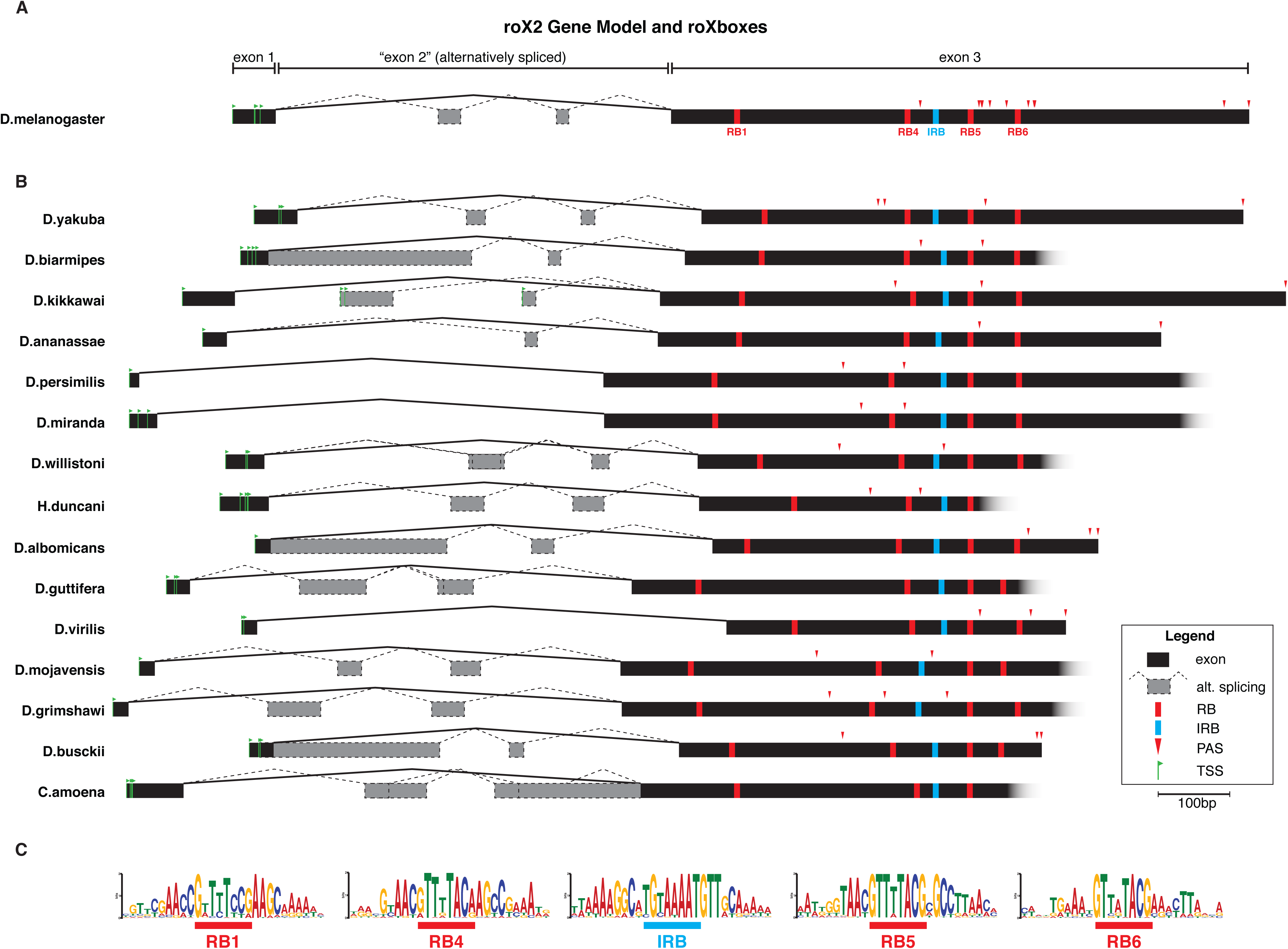
roX2 orthologs have similar gene model, roXbox content, and roXbox sequences. (A) We performed 5’- and 3’-RACE (rapid amplification of cDNA ends) on a selection of 16 diverse roX2 orthologs to define their gene structure, including transcriptional start sites (TSS; green flags), alternative splicing (gray boxes and dashed lines), and polyadenylation sites (PAS; red arrowheads). These 16 species were selected for their diversity and representation of major fly clades. *D.melanogaster* roX2 is known to have numerous alternative splice forms^61^. The major isoform of *D.melanogaster* roX2 consists of a short first exon (∼30bp), an intron (∼600bp, called “exon 2” despite being an intron), and exon 3 (∼500bp, contains roXboxes and described secondary structures); however, many minor isoforms are generated by alternative splicing within “exon 2”. (B) RACE showed that all orthologs analyzed share a similar gene structure: a major isoform consisting of exon 1 – exon 3 with minor isoforms from alternative splicing of “exon 2”. TSS within the short first exon vary, as do PAS within exon 3, occurring most commonly 3’ of roXbox-4, -5, or -6. The relative positions, number, and orientation of roXboxes (RB, red blocks) and inverted roXboxes (IRB, cyan blocks) are consistent across most species. Here, graphical alignment is relative to RB5, the most highly conserved of these sequences. (C) Indeed, the RBs and IRB are the most highly conserved sequence elements in roX2 orthologs. The 8-nucleotide RB or IRB motif is underlined. Motifs were calculated over all discovered roX2 (and roX3) orthologs. Other weakly conserved sequences occur within “exon 2” and are likely implicated in alternative splicing (not shown).

**Figure Supplement 4.**
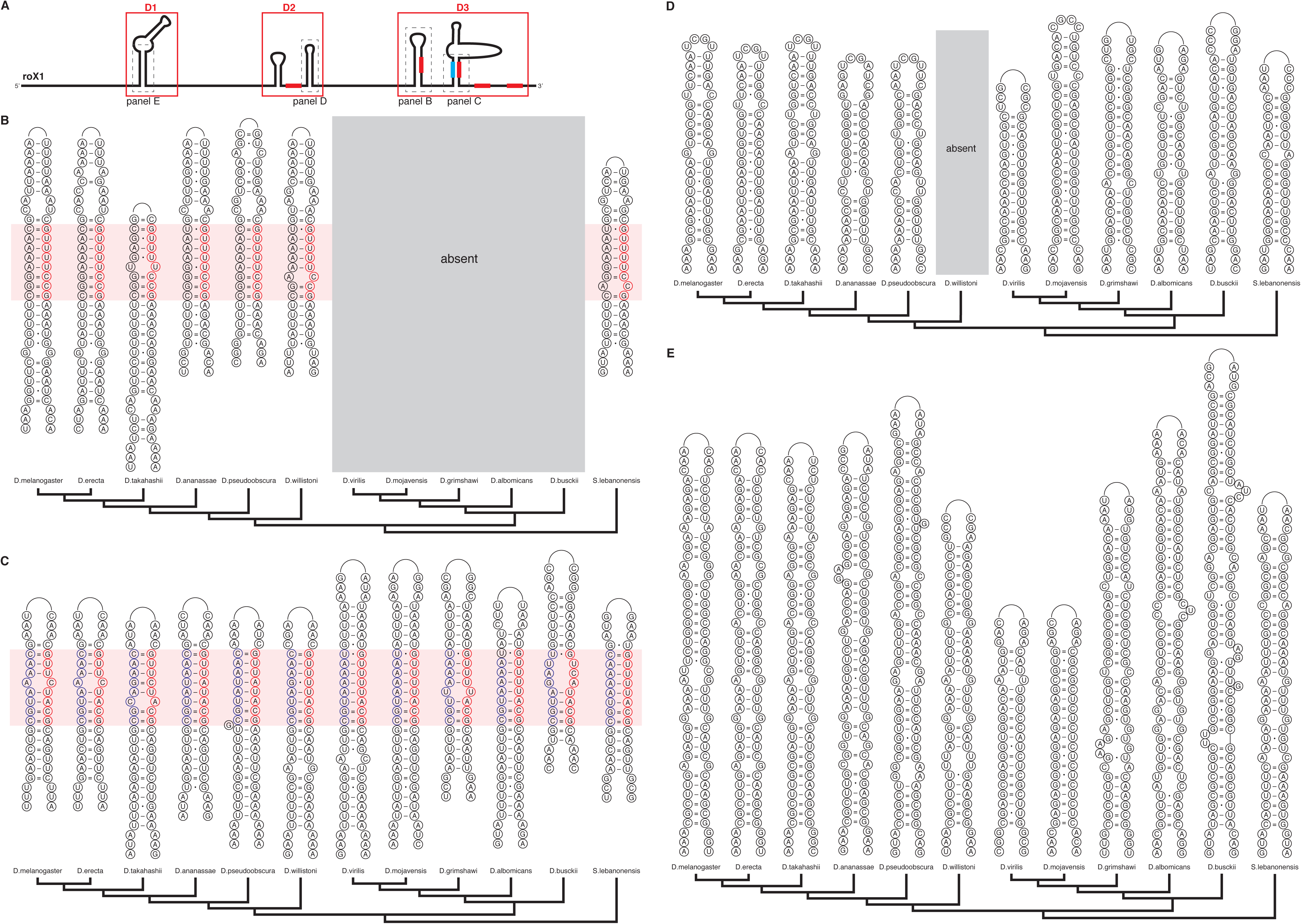
roX1 orthologs have similar secondary structures, but the Drosophila subgenus lacks a critical stem-loop in domain D3 and a structure in D2 is lost in *D.willistoni*. (A) The secondary structures of roX1 in *D. melanogaster* are organized around three primary functional domains: D1, D2, and D3 (red boxes). D3 contains four RBs and one IRB (red and cyan blocks, respectively), and D2 contains one RB. The indicated structures and their conservation are shown in panels **B-E**. All structures are drawn 5’-to-3’, left-to-right. For brevity, only representative species from major clades are shown. (B) The stem-loop within D3 of roX1 contains a RB1 (red highlight) and is conserved in the Sophophora subgenus, but both the roXbox and the structure are absent in the Drosophila subgenus. The stem-loop and RB1 are present in the outgroup species, *S.lebanonensis* See also **Figure 3C-D**. (C) The IRB-RB2 (red highlight) stem-loop within D3 of roX1 is conserved across all *Drosophila* species. (D) The stem-loops within domain D2 are absent in *D.willistoni*, which has no apparent D2 domain. Most species in the Drosophila subgenus also lack the first stem-loop structure in D2 (not shown). (E) The primary stem of domain D1 is found in all species, and is proximal to the MRE within the *roX1* locus (which acts as an active HAS in all species investigated).

**Figure Supplement 5.**
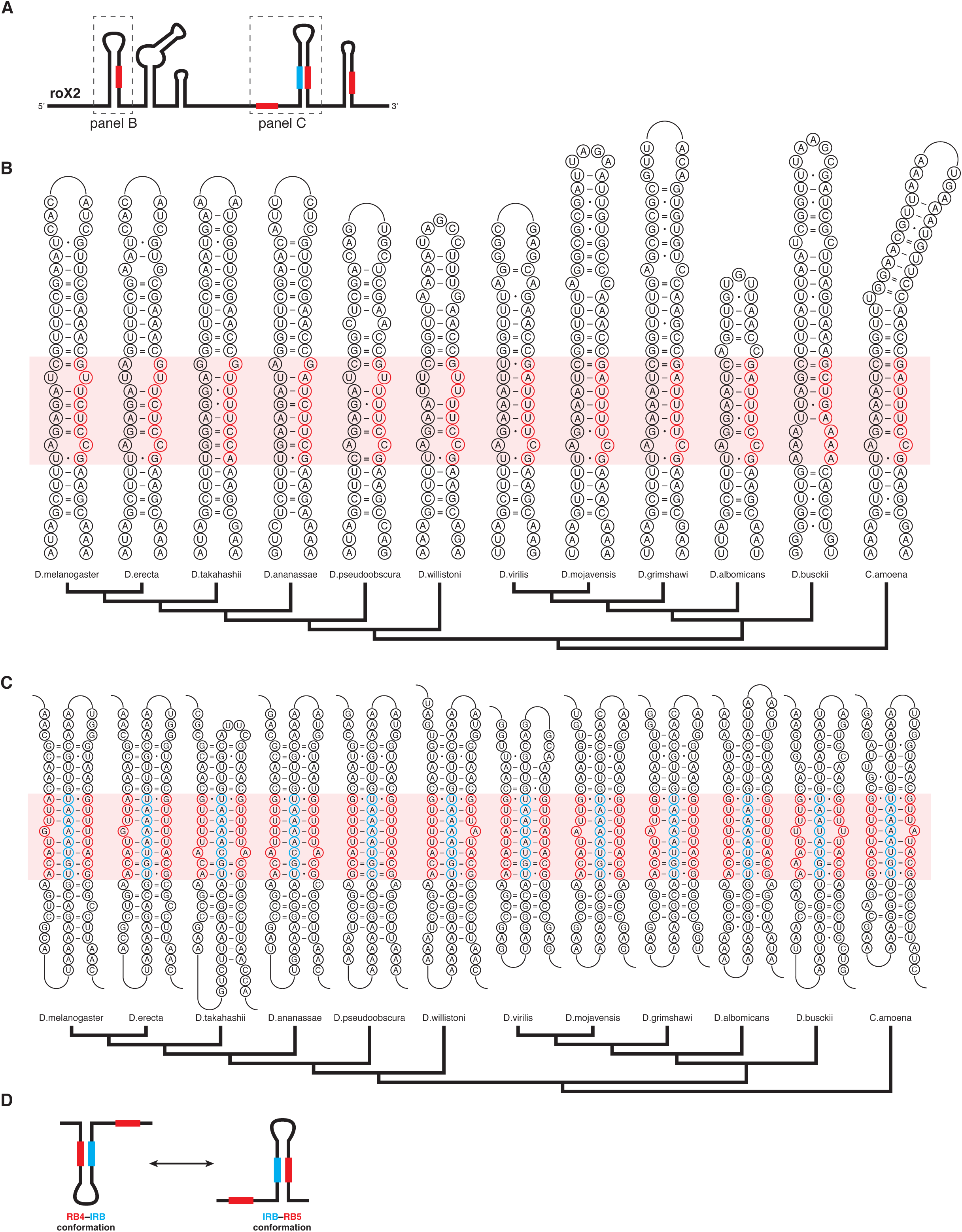
roX2 orthologs have similar secondary structures, including a conserved pair of alternative secondary structures between RB4, IRB, and RB5. (A) The structure of model *D.melanogaster* roX2, all within exon 3. The indicated structures and their conservation are shown in panels **B-C**. All structures are drawn 5’-to-3’, left-to-right. For brevity, only representative species from major clades are shown. (B) The stem-loop at the 5’-end of roX2 contains a roXbox (RB1) and is conserved across all Drosophila species. (C-D) Two alternative structures are ultraconserved across every species, and involve RB4, IRB, and RB5. RB4 and RB5 compete for the single intervening IRB to form two alternative and mutually exclusive structures, the RB4–IRB conformation or the IRB–RB5 conformation. Both structural forms are shown simultaneously here, competing for the central IRB element, in cyan (note: structure cartoon does *not* depict an RNA triplex).

**Figure Supplement 6.**
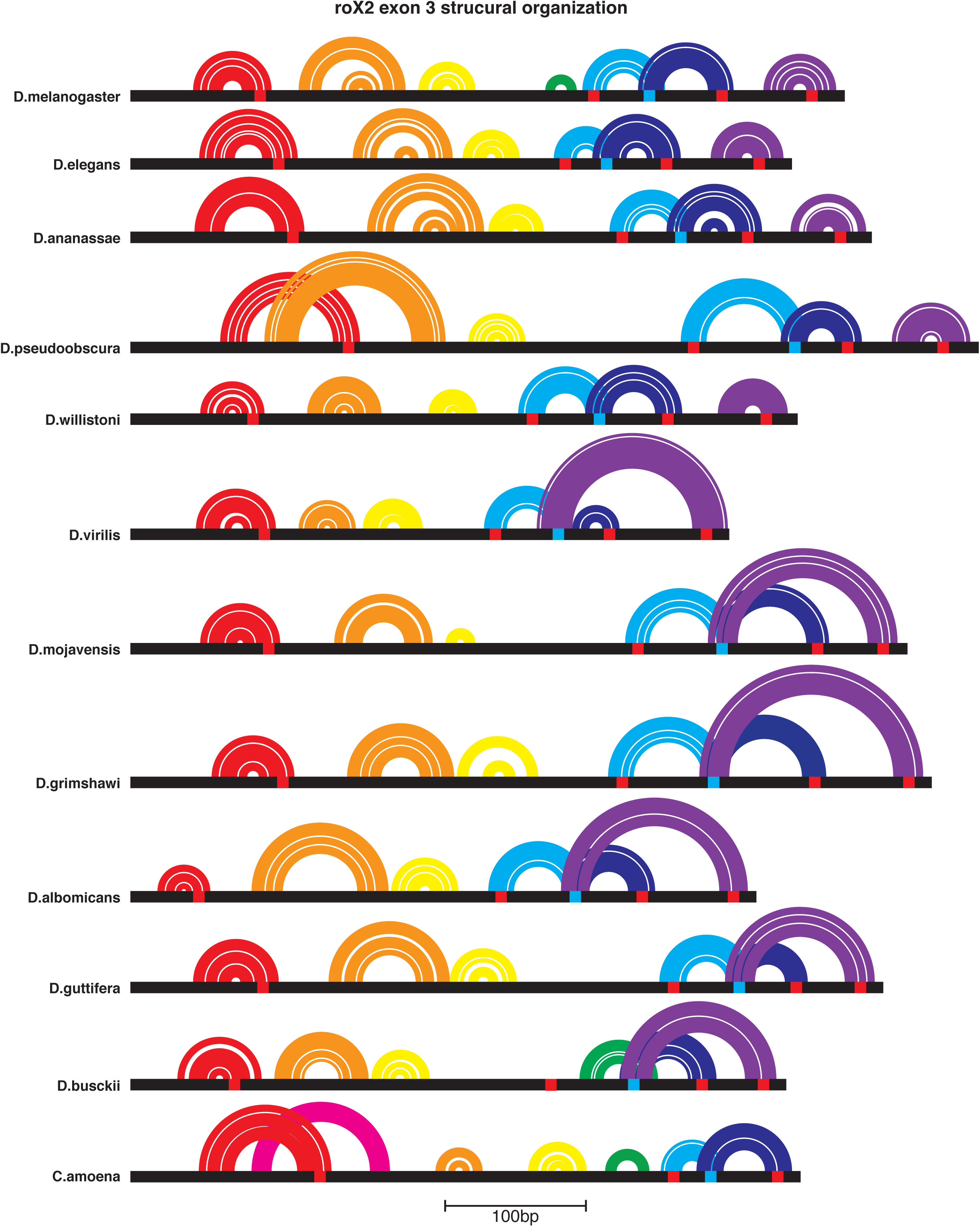
The structural organization of roX2 exon 3 is conserved. RB (red blocks) and IRB (cyan blocks) fold into stem-loops (colored arcs; circle plot). The P1 stem-loop (red arcs) and RB4–IRB–RB5 alternative structures (cyan, blue, and purple) are the most conserved structures; see also **figure supplement 5**. For brevity, only representative species from major clades are shown.

**Figure Supplement 7.**
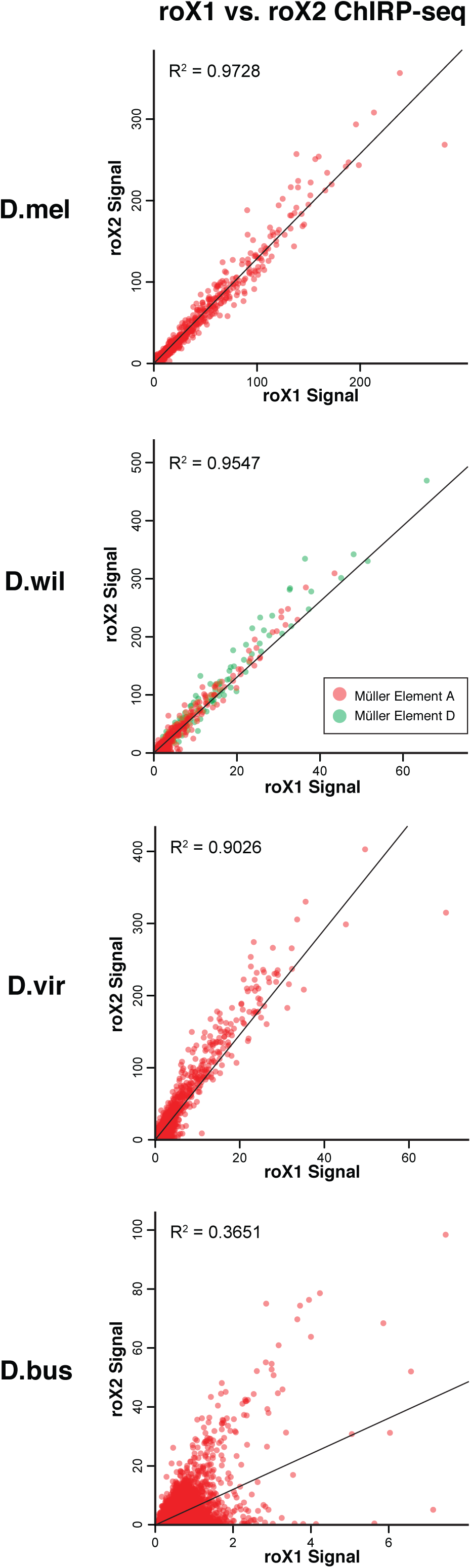
Within each species, roX1 and roX2 have the same binding sites, though with different absolute affinities. In all species, roX1 and roX2 bind to the same loci, though some are biased towards roX2. The signal from roX1 or roX2 ChIRP-seq in 1kb windows of the X-chromosome (ME-A in all species, plus ME- D in *D.willistoni*) was integrated, and plotted against one another. There is high correlation between roX1 and roX2 signal, especially for *D.melanogaster, D.willistoni*, and *D.virilis* with greatly diminished correlation in *D.busckii*. The roX1 signal is substantially lower than roX2 signal for all species except *D.melanogaster*, indicating a bias towards roX2 as the dominant roX homolog in these species (see **Figure 3C**; note that *x-* and *y-* axes are not equally scaled). The 5kb windows surrounding the *roX1* and *roX2* loci were excluded due to direct ChIRP oligo–genomic DNA hybridization and recovery.

**Figure Supplement 8.**
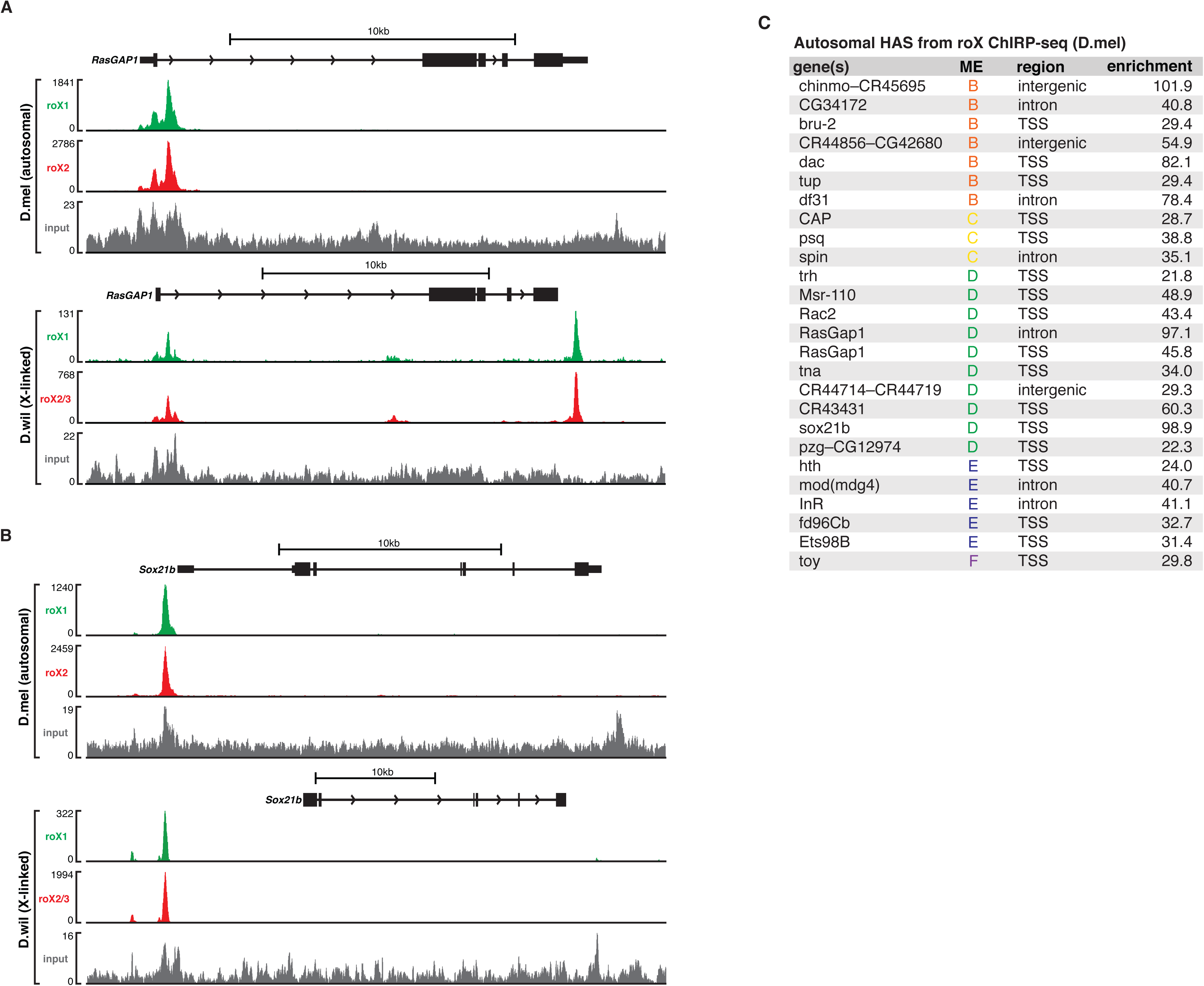
roX RNAs bind to dozens of autosomal sites, some of which are conserved X-linked HAS in *D.willistoni*. (A-B) Two such HAS at the TSS / promoters of autosomal genes (on ME-D) are shown, *RasGAP1* and *Sox21b*. These genes are autosomal in *D.melanogaster* (on ME-D), but X-linked in *D.willistoni* (due to its ME-A+D fusion). The HAS at the TSS / promoters of these two genes are conserved between *D.melanogaster* and *D.willistoni*; however, in *D.willistoni* an additional HAS is immediately downstream from the *RasGAP1* stop codon (presumably in the 3’UTR). This suggests that preexisting autosomal binding sites may also serve as HAS after neo-sex chromosome karyotype fusions. (C) The autosomal HAS in *D.melanogaster* are present on all autosomes (ME-B through -F), and are predominantly at TSS / promoters (as opposed to the intronic and 3’-UTR bias of X-linked HAS). These autosomal HAS are reproducible across roX1 and roX2 ChIRP-seq and between different ChIRP-seq experiments in different cell types^37^. Interestingly, some of these genes have male-specific or male-biased expression, such as *chinmo*, *Sox21b*, and *dac* (not shown), suggesting that male-biased autosomal genes may coopt the dosage compensation complex to upregulate expression in a male-specific manner.

**Figure Supplement 9.**
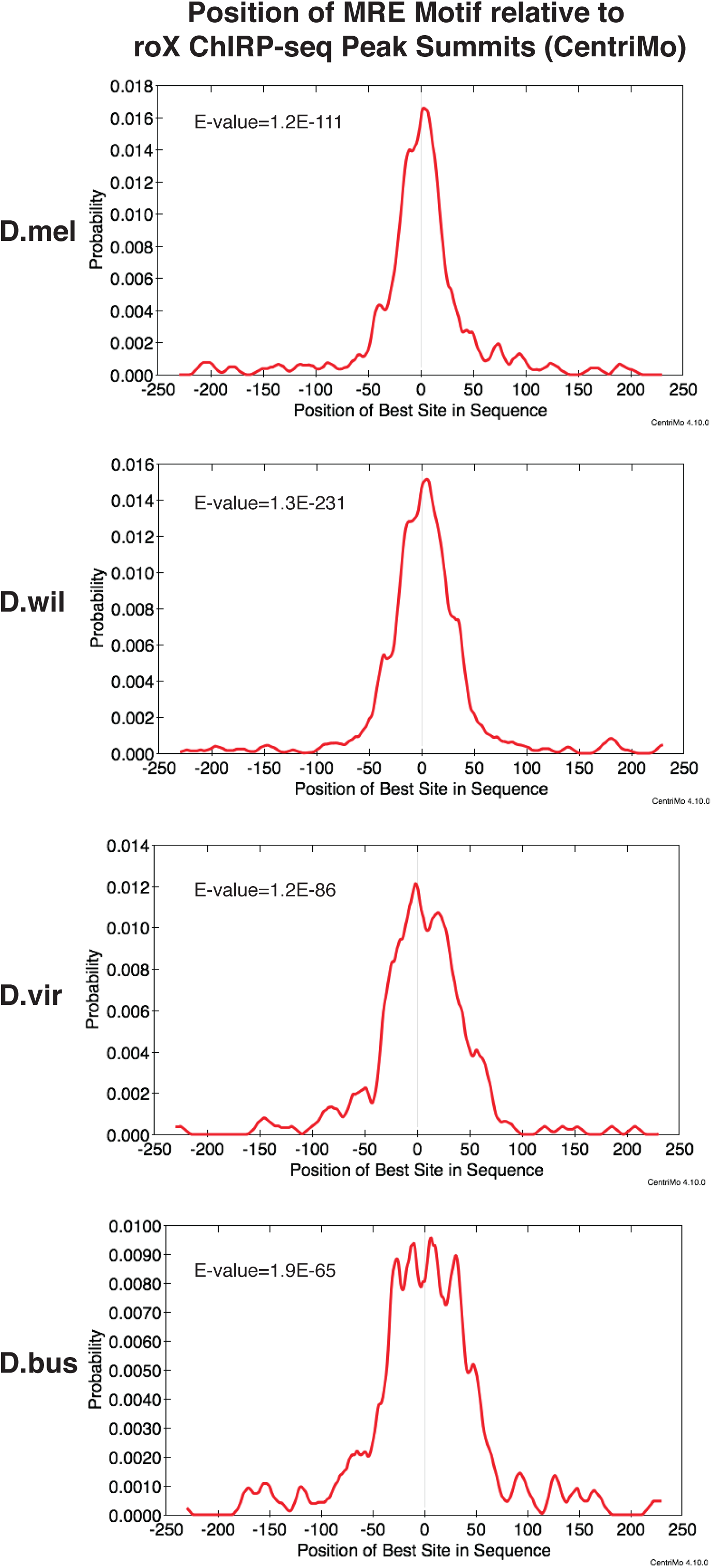
The MRE motif is centered at roX2 ChIRP-seq peaks (HAS). The best-matched MRE motif within each roX2 ChIRP-seq peak (HAS) was calculated by CentriMo^50^ and plotted. In each species, the MRE motif is significantly centered, indicating the precision and high-fidelity with which ChIRP-seq can map precise roX binding sites.

**Figure Supplement 10.**
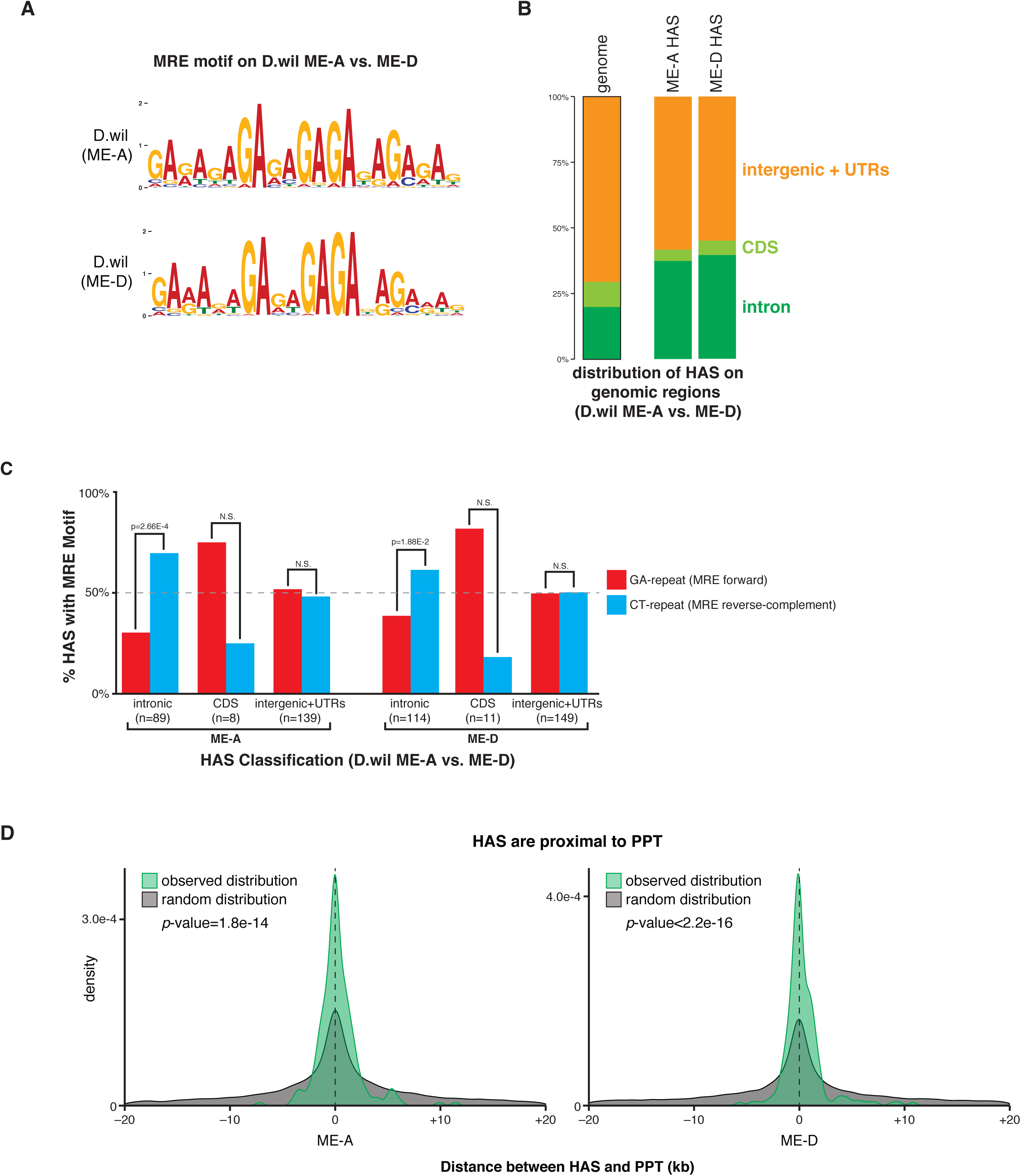
HAS on *D.willistoni* ME-D are similar to HAS on ME-A, despite its ancestry as an autosome. (A) The MRE motifs from ME-A and ME-D are indistinguishable. We did not find evidence of tamed transposable elements at HAS on ME-D, as found on the more recently evolved neo-sex chromosome of the *D.miranda*^40^. (B) ME-A and ME-D HAS are similarly distributed on genomic regions, and are enriched especially in introns. Note that UTRs are grouped with intergenic regions here. (C) Intronic HAS are significantly biased towards the reverse-complement orientation of the MRE motif (CT-repeat) on both ME-A and ME-D. (D) HAS are significantly proximal to PPT on both ME-A and ME-D.

**Figure Supplement 11.**
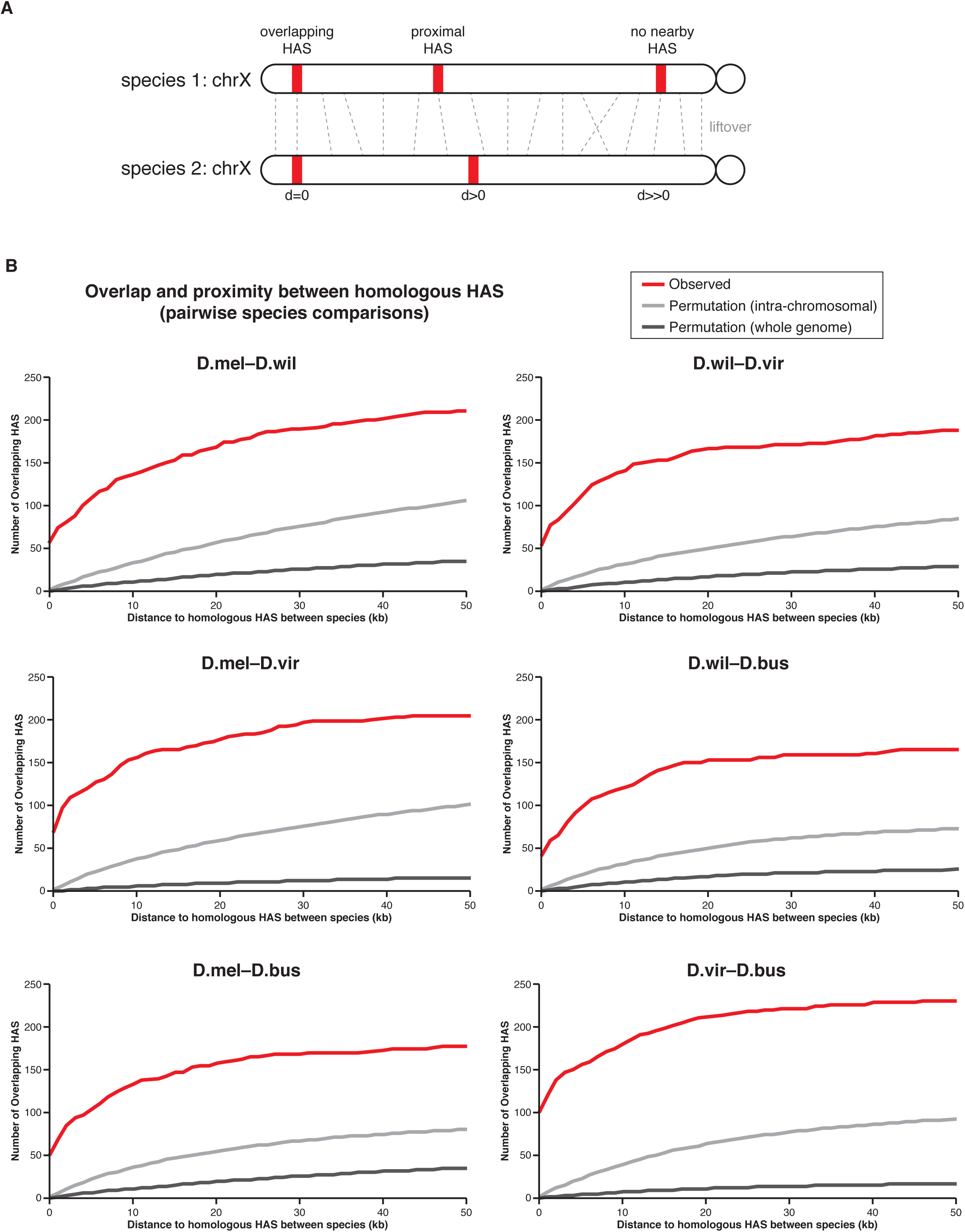
Overlap and proximity between homologous roX binding sites. (A) Species-to-species liftover and HAS distance calculation strategy. Homologous regions in two species’ genomes were mapped by genome-wide liftover. If homologous sites are both HAS, the HAS are overlapping and the distance between is 0. If a homologous site is a HAS in one species and not another, the distance to the nearest HAS is calculated (d>0). If the HAS has no nearby neighbor in the other species, the distance is much larger. (B) Overlap and proximity between homologous HAS using the above strategy for all pairwise species comparisons. Though exact conservation of binding sites (i.e. distance = 0) between any two species is limited (approximately 10-30%), this is significantly higher than expected by random chance, as a random permutation of all HAS over their respective chromosomes or the whole genome yields very few overlapping peaks. Additionally, if exact peak overlap is lost (i.e. d>0), there is a high likelihood that another peak is nearby in the homologous genomic region, as indicated by the steep slope of the “observed” line in this distance regime. A higher percentage of peaks overlap in D.mel–D.wil and D.vir–D.bus than in other pairwise comparisons, perhaps reflecting the closer phylogenetic relationships of these pairs.

**Figure Supplement 12.**
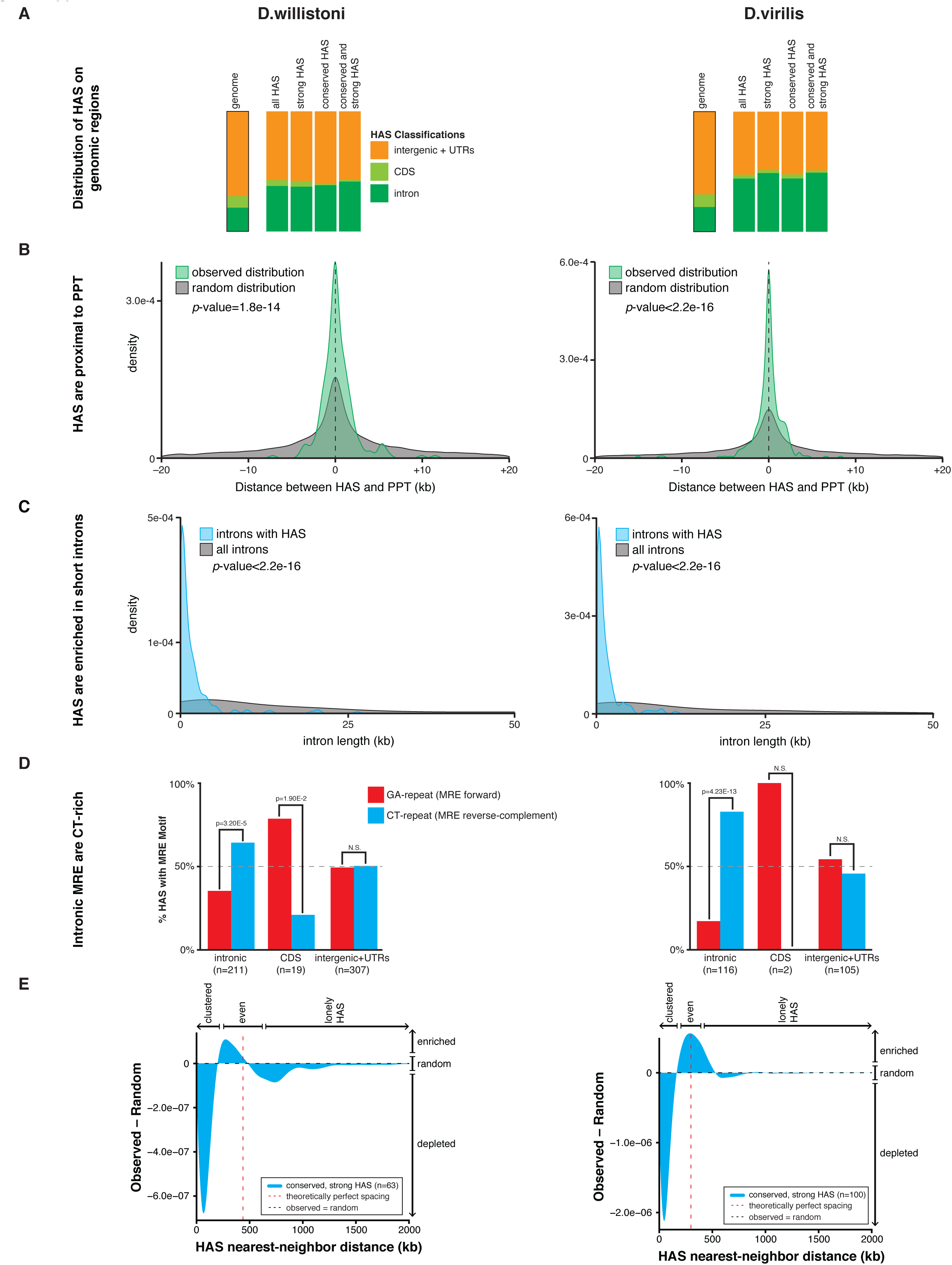
Analyses of HAS in *D.willistoni* and *D.virilis* (supplementary to *D.melanogaster* analyses in Fig. 5). (A) HAS are distributed over genomic regions (intergenic + UTRs, CDS, and introns), with particular enrichment in introns. Intergenic and UTRs are grouped together because UTRs are not reliably mapped in these species. (B) HAS are significantly proximal to PPT in both *D.willistoni* and *D.virilis*. (C) HAS are enriched in short introns in both *D.willistoni* and *D.virilis*, relative to all introns. (D) Intronic HAS are significantly biased towards the reverse-complement orientation of the MRE motif (CT-repeat) in both *D.willistoni* and *D.virilis*. There is a weak bias towards the GA-repeat MRE motif in CDS in *D.willistoni*. Note that peak classifications differ from those in **Figure 5**. (E) The difference between the observed and random HAS (conserved, strong only) distributions on the X-chromosomes of *D.willistoni* and *D.virilis*. The positive *y*-value near the theoretically perfect spacing distance indicates an enrichment of the even spacing model relative to random spacing; conversely, the negative y-value at short distances indicates a depletion of the clustered spacing model relative to random spacing. This trend is not as robust in *D.willistoni*.

**Figure Supplement 13.**
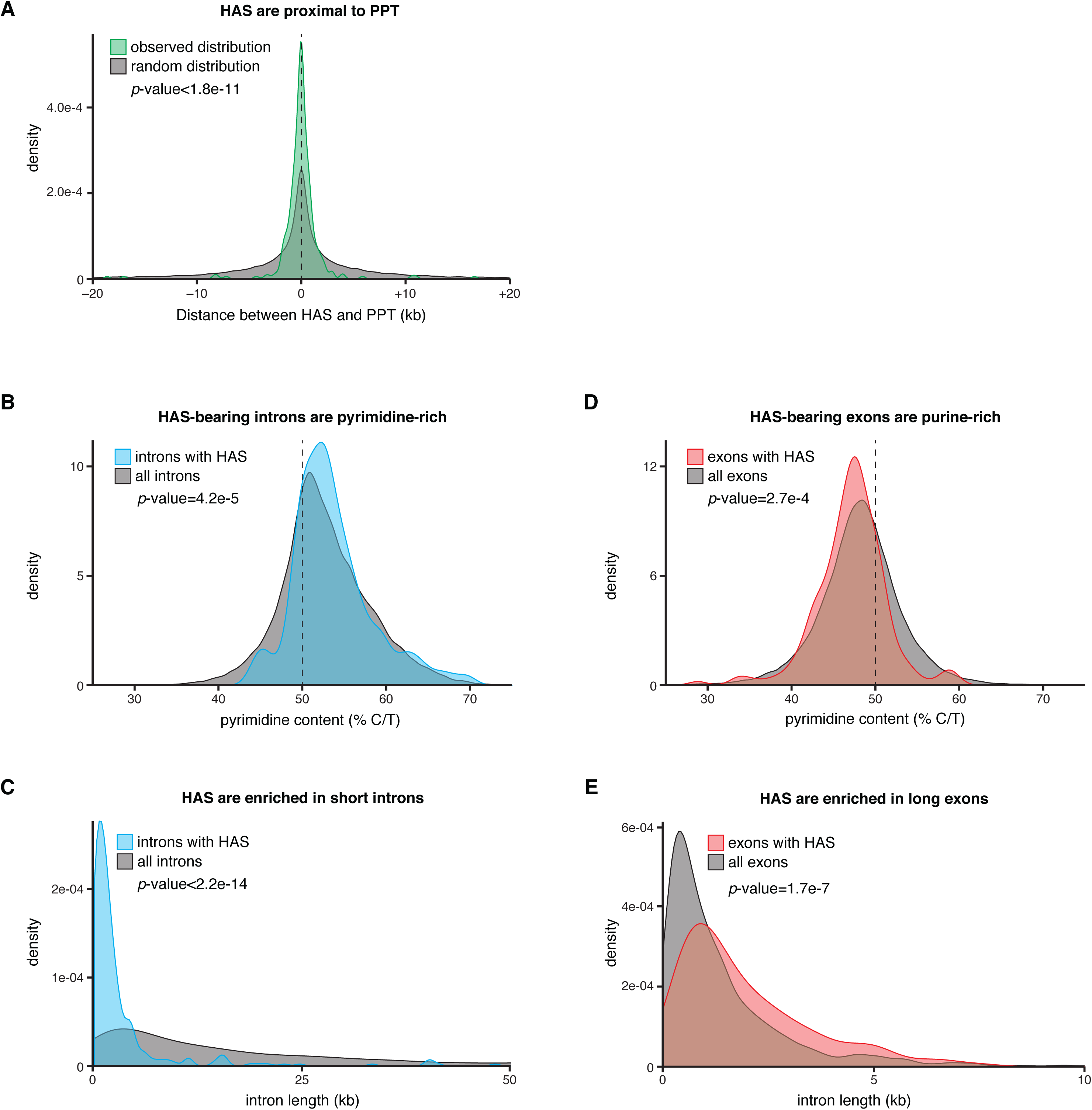
Analysis of intronic and exonic HAS in *D.melanogaster*. (A) HAS are significantly proximal to PPT relative to a random distribution. Approximately 20% of HAS are ±100bp from a PPT, vs. 7% in the random distribution. (B) HAS-bearing introns are more pyrimidine-rich than typical introns. (C) HAS-bearing introns are shorter than typical introns. (D) HAS-bearing exons are more purine-rich than typical exons. (E) HAS-bearing exons are slightly longer than typical exons.

**Figure Supplement 14.**
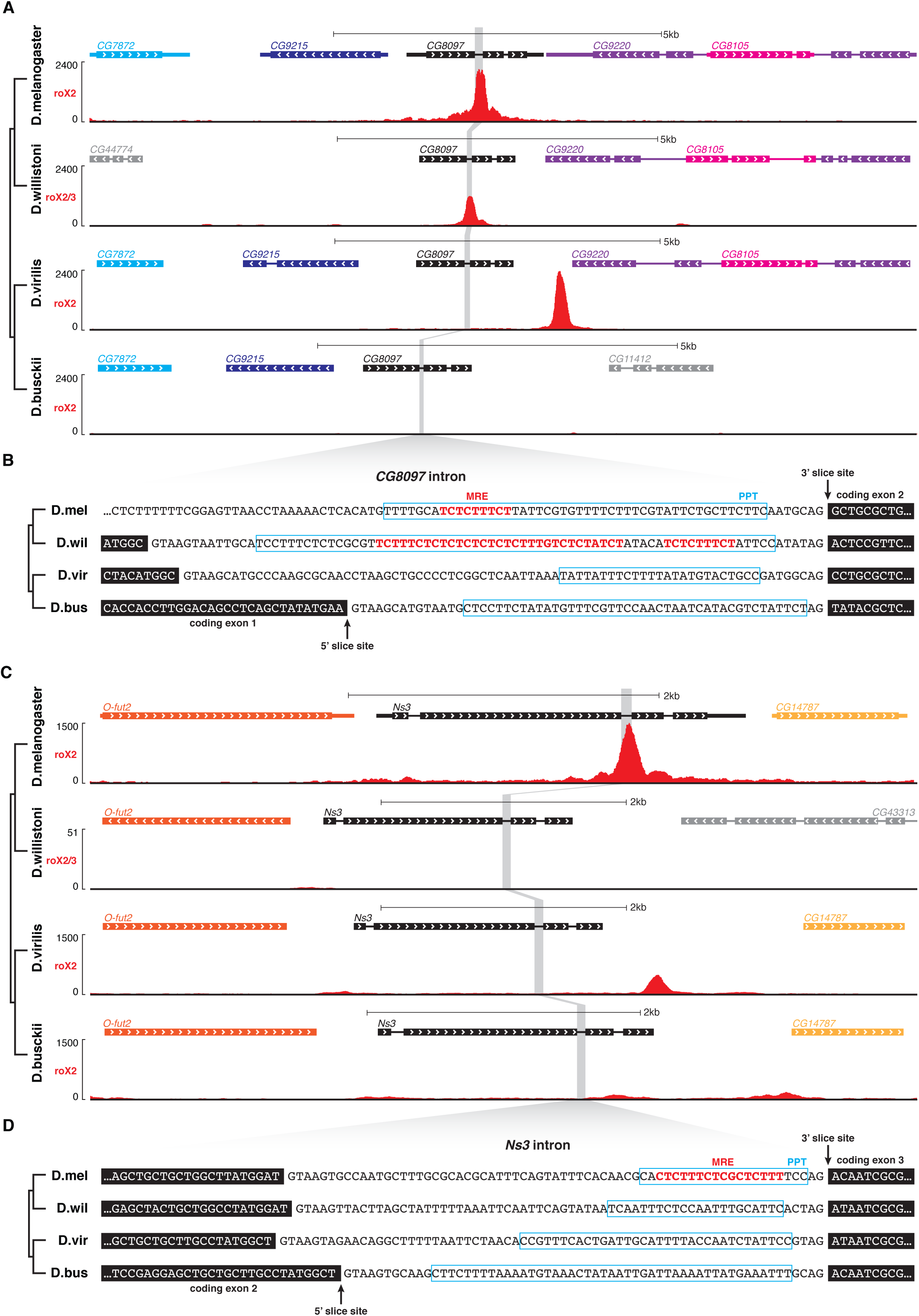
MRE can evolve from the polypyrimidine tract of introns. (A) Genome browser tracks of roX2 ChIRP-seq at the *CG8097* locus in four species. In *D.melanogaster* and *D.willistoni*, there is a HAS in the first intron of *CG8097* (highlighted in gray), but this HAS is lost in *D.virilis* and *D.busckii*. In *D.virilis*, a new HAS is present in the putative 3’UTR of neighboring gene *CG9220*, illustrating the principle of HAS turnover in proximity. (B) The highlighted sequence of the first intron of *CG8097* in four species (coding exons shown as black blocks). Instances of the MRE motif (red) are present within the pyrimidine-rich PPT (cyan box, approximately) of *CG8097*’s intron in *D.melanogaster* and *D.willistoni.* However, high-scoring incidences of the MRE motif are absent from the PPT of *CG8097*’s intron in *D.virilis* and *D.busckii*, in accord with the lack of a corresponding HAS in these two species. Thus, intronic PPT can serve as both an RNA signal (for splicing) and as a DCC binding site (MRE-bearing HAS). (C) Genome browser tracks of roX ChIRP-seq at the *Ns3* locus in four species. In *D.melanogaster*, there is a HAS in the second intron of *Ns3* (highlighted in gray), but this HAS is absent in *D.willistoni*, *D.virilis,* and *D.busckii*. Most parsimoniously, this suggests that the HAS evolved in the *melanogaster* lineage. In *D.virilis*, a new HAS is present nearby, illustrating the principle of HAS turnover in proximity. (D) The highlighted sequence of the second intron of *Ns3* in four species. Again, the MRE motif is present within the PPT of *Ns3’s* intron in *D.melanogaster* and no high-scoring MRE motifs are found in *D.willistoni*, *D.virilis,* and *D.busckii*, in accord with the lack of a corresponding HAS in these three species.

**Figure Supplement 15.**
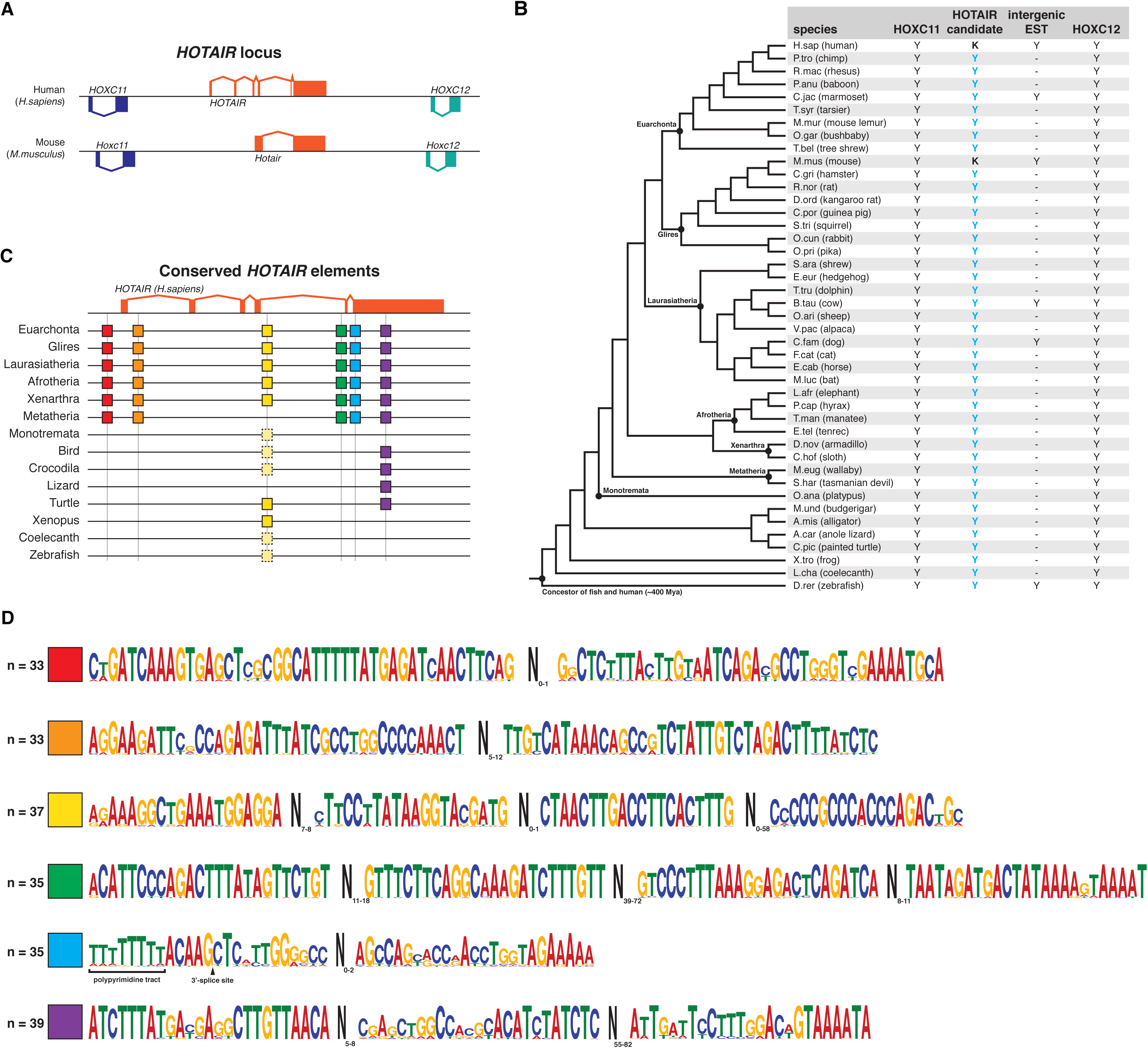
The lncRNA ortholog search strategy found the *HOTAIR* locus in 43 diverse vertebrate species. (A) HOTAIR is a lncRNA that was discovered in human and has been described in mouse. It is transcribed from the *HOXC* locus, flanked by and antisense to the protein-coding genes *HOXC11* and *HOXC12*, which are highly conserved across vertebrate genomes. (B) We searched for the *HOTAIR* locus in 43 diverse vertebrate species from primates down to zebrafish (which diverged ∼400Mya). We used the synteny module of the lncRNA ortholog search strategy, initiating with knowledge of only human HOTAIR. We found *HOXC11* and *HOXC12* on the same genomic scaffold in a window of ∼21kb, suggesting that the syntenic relationship with *HOTAIR* is maintained. In at least six species for which there are expressed sequencing tags (ESTs), there was an EST in the intergenic space between *HOXC11* and *HOXC12*, mapping to the location where *HOTAIR* would be expected. This suggests that an intergenic lncRNA – presumably the HOTAIR ortholog – is encoded at this locus. K, known HOTAIR lncRNA; Y (cyan), HOTAIR lncRNA ortholog candidate identified. (C) Using the motif discovery algorithm MEME, we searched for instances of microhomology in the putative HOTAIR loci. At least six instances of focal microhomology were found, represented as colored boxes and corresponding to specific positions mapped to human HOTAIR. All six elements (red through purple) are conserved across all eutherian mammals, while one (yellow) has much deeper evolutionary conservation and can be found in zebrafish. Despite monotremes’ closer relation to eutherian and metatherian mammals, many of the sequence elements found in HOTAIR are absent. Pale boxes with dotted outline indicate species for which not all elements within a microhomologous sequence motif are present (i.e. incomplete microhomology). (D) Sequence motifs for the conserved HOTAIR elements. One motif (red) is found at the promoter for the human HOTAIR lncRNA, and its conservation suggests that this promoter is conserved in eutherian and metatherian mammals. Similarly, a splice site (cyan) is conserved in eutherian and metatherian mammals. Together, the conservation of these transcription- and splicing-associated signals suggests that this locus in other species is also transcribed and spliced. Whether these are microhomologous elements are functional at the DNA level (e.g. transcription factor binding sites, enhancers, etc.) or at the RNA level (RNA-binding protein sites, RNA processing sites, microRNA targets, etc.) and their importance to HOTAIR function remains to be validated.

